# Metagenomic assemblies tend to break around antibiotic resistance genes

**DOI:** 10.1101/2023.12.13.571436

**Authors:** Anna Abramova, Antti Karkman, Johan Bengtsson-Palme

**Affiliations:** Department of Infectious Diseases, Institute of Biomedicine, The Sahlgrenska Academy, University of Gothenburg, Guldhedsgatan 10A, SE-413 46 Gothenburg, Sweden; Division of Systems and Synthetic Biology, Department of Life Sciences, SciLifeLab, Chalmers University of Technology, SE-412 96 Gothenburg, Sweden; Centre for Antibiotic Resistance research (CARe) in Gothenburg, Sweden; Department of Microbiology, University of Helsinki, Helsinki, Finland

**Keywords:** metagenomic assembly, antibiotic resistance genes, genomic context

## Abstract

**Background:** Assembly of metagenomic samples can provide essential information about the mobility potential and taxonomic origin of antibiotic resistance genes (ARGs), and inform interventions to prevent further spread of resistant bacteria. However, ARGs typically occur in multiple genomic contexts across different species, representing a considerable challenge for the assembly process. Usually, this results in many fragmented contigs of unclear origin, complicating the risk assessment of ARG detections. To systematically investigate the impact of this issue on detection, quantification and contextualization of ARGs, we evaluated the performance of different assembly approaches, including genomic-, metagenomic- and transcriptomic-specialized assemblers. We quantified recovery and accuracy rates of each tool for ARGs both from *in silico* spiked metagenomic samples as well as real samples sequenced using both long- and short-read sequencing technologies.

**Results:** The results revealed that none of the investigated tools can accurately capture genomic contexts present in samples of high complexity. The transcriptomic assembler Trinity showed a better performance in terms of reconstructing longer and fewer contigs matching unique genomic contexts, which can be beneficial for deciphering the taxonomic origin of ARGs. The currently commonly used metagenomic assembly tools metaSPAdes and MEGAHIT were able to identify the ARG repertoire but failed to fully recover the diversity of genomic contexts present in a sample. On top of that, in a complex scenario MEGAHIT produced very short contigs, which can lead to considerable underestimation of the resistome in a given sample.

**Conclusions:** Our study shows that metaSPAdes and Trinity would be the preferable tools in terms of accuracy to recover correct genomic contexts around ARGs in metagenomic samples characterized by uneven coverages. Overall, the inability of assemblers to reconstruct long ARG-containing contigs has impacts on ARG quantification, suggesting that directly mapping reads to an ARG database might be necessary, at least as a complementary strategy, to get accurate ARG abundance and diversity measures.

## Background

Antimicrobial resistance (AMR) is an increasing global health crisis causing hundreds of thousands of deaths each year worldwide (<u>Murray et al. 2022</u>). To limit its spread, there is a need to identify and quantify resistance in both clinical and environmental settings. Metagenomic sequencing is a powerful tool allowing simultaneous identification and quantification of antibiotic resistance genes (ARGs) in a given sample. Metagenomic analysis of sewage from different parts of the world has revealed that the same ARGs are found in different genomic backgrounds globally, proving the need to not only identify the composition of ARGs in a given sample, but also in what genomic context they are present. The genomic background of an ARG determines co-resistance patterns and mobilization potential, both of which can affect the choice of intervention strategies locally and globally (<u>Munk et al. 2022</u>). For this reason, metagenomic sequencing has been suggested as a possible means for surveillance of AMR not only in sewage (<u>Hendriksen et al. 2019</u>; <u>Pruden et al. 2021</u>), but also in the environment in general (<u>Bengtsson-Palme et al. 2023</u>). Current high-throughput sequencing platforms produce hundreds of millions of reads that require assembly to be reconstructed into longer stretches called contigs, which can provide more contextual information. This step is typically demanding in terms of computational resources and time (<u>Vollmers et al. 2017</u>; <u>Meyer et al. 2022</u>). On top of this, short read length, skewed species abundance distributions, high similarity between closely related ARG variants, and massive amounts of data make recovery of ARGs and the context around them challenging from metagenomic data (<u>Ayling et al. 2019</u>; <u>Yorki et al. 2023</u>).

There are currently several tools available to assemble short-read sequencing data from metagenomic samples (see review by <u>Ayling et al. (2019)</u>), most of which use variants of the de Bruijn graph approach to handle large amounts of data in an efficient way. This approach is based on reconstructing graphs to represent k-mers present in a set of reads, followed by traversing these graphs and identifying the most probable path representing a contig. Converting a graph path into a contig is not a trivial task. Metagenomic samples typically contain an unknown number of species with unknown abundance distributions. In the case of related species, sequences can carry similar sets of k-mers resulting in complex assembly graphs. This is further complicated by conserved repetitive regions (such as ARGs or ribosomal RNA genes). Assembling conserved regions present in several different genomic contexts typically results in highly complex branched assembly graphs, which makes traversing the graphs extremely difficult. This is generally solved by splitting the graph into multiple short contigs (<u>Bengtsson-Palme et al. 2017</u>). For metagenomic analysis targeting ARGs, this means that sometimes all contextual information regarding the taxonomic origin or mobility of a gene will be lost, which can potentially lead to misinterpretation of the results.

There are several studies benchmarking metagenomic assembly tools, such as the “Critical Assessment of Metagenome Interpretation” (CAMI) challenge (<u>Wang et al. 2019</u>; <u>Meyer et al. 2022</u>). The focus of these studies has largely been on the ability of assemblers to distinguish evolutionary related organisms in complex microbial samples. There is also a study by <u>Brown et al. (2021)</u> assessing different assemblers for contextualization of ARGs using co-occurrence of ARGs and mobile genetic elements (MGEs) on assembled contigs as a proxy. However, a critical evaluation of currently available short-read assemblers on reconstructing the context around ARGs existing in multiple genomic contexts is currently lacking. ARGs constitute a type of genomic feature that is particularly likely to be fragmented in metagenomic assemblies, as they are often present in multiple contexts, can be surrounded by various forms of repeat regions, and can be present on plasmids with varying degrees of copy numbers. A specific investigation of how assemblers handle these genes is therefore warranted. Furthermore, the resulting assemblies are often used to perform ARG quantification by mapping reads back to the contigs to estimate gene abundances. It is not clear how the choice of assembler will affect this form of ARG quantification and, by consequence, the final biological interpretation of the results.

The main goal of this study was to systematically evaluate the capability of assembly tools to recover ARGs in the correct genomic context from metagenomic data. To have a controlled but still real-life relevant experimental set up, we first used a real data set from human stool samples and spiked it with simulated reads derived from resistance plasmids. We then assembled the test datasets and evaluated performance of several tools (Velvet, SPAdes, metaSPAdes, MEGAHIT, Trinity, Ray) with respect to their accuracy of recovering the genomic contexts of ARGs using the original plasmids as reference. Furthermore, we did the same assessment but on a sample sequenced with both short- and long-read technologies, using the latter as a reference. The results provide important perspectives on the choice of assembly programs for recovering correct genomic contexts for ARGs from metagenomic samples. Furthermore, they call into question some of the practices currently used for quantification of ARGs based on metagenomic sequencing.

## Methods

### Evaluation using simulated reads

To obtain a controlled experimental setup, we randomly selected a metagenomic dataset and spiked it with simulated reads derived from plasmids containing a known set of ARGs. The metagenomic dataset was downloaded from Sequence Read Archive (SRA) and corresponds to a human stool sample, representing a common sample type used for studies of AMR. This sample was sequenced by Illumina NextSeq550 with 150 bp reads, resulting in 4.1Gb dataset (SRR9654970). To obtain a set of plasmids, we first chose a number of clinically-relevant and commonly observed ARGs from different ARG classes, including *sul2* (816 bp), *blaNDM-1* (813 bp), *blaTEM* (861 bp), *aph(3’’)-Ib_3* (804 bp) and *tet(A)* (1200 bp). We downloaded protein sequences from the Comprehensive Antibiotic Resistance Database (CARD) database and used them as queries for NCBI BLAST database searches to retrieve complete plasmid sequences. Only hits with >98% identity to the ARG query and corresponding to full-length plasmids were selected, five for each selected ARG (Table 1). We aimed to select plasmids of different sizes to reflect the diversity in natural samples. For this test, the number of plasmids was selected arbitrarily, but it is comparative to real sample complexity; e.g. 26 known and 21 putative novel plasmids were recovered in an Indian lake metagenome (<u>Bengtsson-Palme et al. 2014</u>).

**Table 1.** Set of plasmids and corresponding ARGs.

| ARG | Accession | Species | Strain | Plasmid | Length, bp | Assigned read coverage proportion |
| --- | --- | --- | --- | --- | --- | --- |
| aph(3'')-Ib_3 | CP039146.1 | *Acinetobacter* sp. | 10FS3-1 | p10FS3-1-3 | 73803 | 5 |
| aph(3'')-Ib_3 | CP058166.1 | *Enterobacter hormaechei* | RHBSTW-00070 | pRHBSTW-00070_3 | 9923 | 34 |
| aph(3'')-Ib_3 | CP026933.2 | *Escherichia coli* | CFS3273 | pCFS3273-1 | 268665 | 1 |
| aph(3'')-Ib_3 | CP055808.1 | *Escherichia fergusonii* | RHB03-C23 | pRHB03-C23_3 | 35135 | 10 |
| aph(3'')-Ib_3 | CP064948.1 | *Pseudomonas fulva* | ZDHY414 | pVIM-24-ZDHY414 | 589460 | 1 |
| blaNDM-1 | AP023079.1 | *Acinetobacter baumannii* | OCU_Ac16a | pOCU_Ac16a_2 | 41087 | 8 |
| blaNDM-1 | CP055250.1 | *Citrobacter freundii* | ZY198 | pZY-NDM1 | 53573 | 6 |
| blaNDM-1 | CP047406.1 | *Escherichia coli* | MS6193 | pMS6193A-NDM | 142890 | 2 |
| blaNDM-1 | CP050380.1 | *Klebsiella pneumoniae* | 51015 | p51015_NDM_1 | 353810 | 1 |
| blaNDM-1 | CP040184.1 | *Raoultella planticola* | Rp_CZ180511 | pRpNDM-1 | 334854 | 1 |
| blaTEM-9 | CP063225.1 | *Enterobacter hormaechei* subsp. *Steigerwaltii* | BD-50-Eh | pBD-50-Eh_2 | 336282 | 1 |
| blaTEM-9 | KR259131.1 | *Escherichia coli* | EC3587 | pEC3587 | 10483 | 32 |
| blaTEM-9 | CP025144.1 | *Klebsiella pneumoniae* | NR5632 | NR5632_p1 | 204123 | 2 |
| blaTEM-9 | GQ160960.2 | *Serratia marcescens* |  | R934 | 13775 | 24 |
| blaTEM-9 | CP024467.1 | *Shigella dysenteriae* | BU53M1 | unnamed1 | 54993 | 6 |
| sul2 | CP059301.1 | *Acinetobacter baumannii* | AC1633 | pAC1633-1 | 174292 | 2 |
| sul2 | CP055707.1 | *Citrobacter freundii* | RHB16-C02 | pRHB16-C02_6 | 6801 | 49 |
| sul2 | CP061493.1 | *Enterobacter hormaechei* subsp. *Xiangfangensis* | GENC284 | pGENC284 | 304958 | 1 |
| sul2 | AP022550.1 | *Escherichia coli* | THO-015 | pTHO-015-1 | 88121 | 4 |
| sul2 | MT415059.1 | *Klebsiella pneumoniae* | NMI3243_13 | pIncR_3243 | 69560 | 5 |
| tet(A)_1 | CP047745.1 | *Enterobacter hormaechei* | Eho-4 | pEcl4-5 | 37460 | 9 |
| tet(A)_1 | AP022535.1 | *Escherichia coli* | THO-006 | pTHO-006-2 | 101966 | 3 |
| tet(A)_1 | CP064130.1 | *Klebsiella pneumoniae* | M911-1 | pM911-1.1 | 75711 | 4 |
| tet(A)_1 | CP065164.1 | *Klebsiella variicola* | KPN029 | unnamed2 | 243621 | 1 |
| tet(A)_1 | CP062224.1 | *Salmonella enterica* subsp. *enterica* serovar *Goldcoast* | R18.1656 | p270k | 270696 | 1 |

We used insilicoseq (<u>Gourlé et al. 2018</u>) to generate simulated reads from plasmids using the NovaSeq error model:

iss generate -g plasmids.fa --abundance_file abundance.txt -m NovaSeq -o reads_file

To fine-tune the distribution of plasmid reads, we provided an abundance.txt file containing proportions of reads weighed according to the size of each plasmid (smaller plasmids get more coverage and larger less, see Table 1, “Assigned read coverage proportion” column). Furthermore, to test how the amount of sequencing data corresponding to plasmids affected the assembly process, we generated read files with different total amounts of simulated reads: 0.5x, 1x, 5x, 10x, where 1x corresponds to the number of reads needed to cover the largest plasmid one time.

To ensure a controlled setup, reads from the human stool dataset were first mapped to the selected set of 25 plasmids and all matches were removed to create a clean test dataset (Figure 1). The cleaned test dataset was then spiked with simulated reads to create four files containing either 0.5x, 1x, 5x or 10x simulated reads each. These four datasets were then assembled by programs using default parameters.

**Figure 1.**
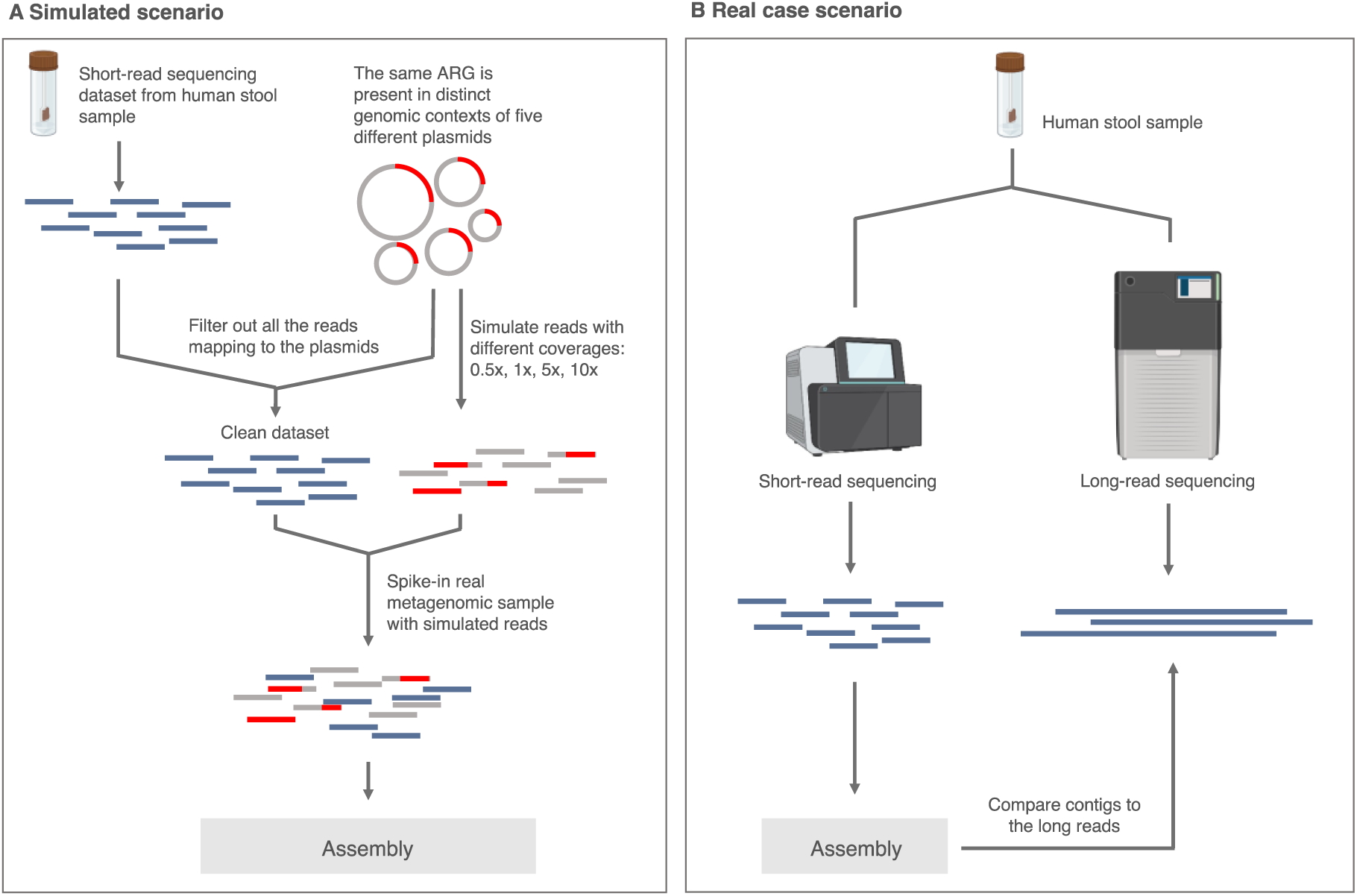
Workflow: A) Simulated scenario constituting a real metagenomic dataset spiked with reads generated from a set of 25 plasmids containing ARGs; B) Real case scenario included long-reads which were used as a reference to quality check the contigs assembled from short read data generated from the same sample.

For this evaluation, we decided to include several different tools: genomic assemblers SPAdes 3.13.0 (<u>Bankevich et al. 2012</u>), Velvet 1.2.10 (<u>Zerbino and Birney 2008</u>) and Ray 2.3.1 (<u>Boisvert et al. 2010</u>); metagenomic assemblers MEGAHIT v1.0.3 (<u>Li et al. 2016</u>) and SPAdes 3.13.0 with the -meta option (also referred to as “metaSPAdes”), and transcriptomic assembler Trinity 2.1.1 (<u>Haas et al. 2013</u>). We also tested TriMetAss as an alternative method to see if it can improve the outcome. TriMetAss is an extension to the Trinity software, which was designed to assemble common and well-conserved genes occurring in multiple genomic contexts in metagenomic data (<u>Bengtsson-Palme et al. 2014</u>).

First, we evaluated assemblers on the selected dataset to assess overall performance. We used METAQUAST 5.2.0 (<u>Mikheenko et al. 2018</u>) to evaluate general assembly performance (Figure S1). Bandage (<u>Wick et al. 2015</u>) tool was used to create assembly graphs visualization.

We also performed the same analyses in a simplified scenario with only two plasmids for each of two resistance genes (*blaNDM-1* and *aph(3’’)-Ib*), to test whether the reduced complexity would improve the assembly performance. We performed this test with CP055250.1 (53573 bp) and AP023079.1 (41087 bp), both carrying the *blaNDM-1* gene, and CP064948.1 (589460 bp) and CP039146.1 (73803 bp) carrying the *aph(3’’)-Ib* gene. To create differential coverages, we generated reads in proportions 2:1 for the first pair and 7:1 for the second pair of plasmids. The reads were generated and spiked into the real dataset as described for the simulated scenario above. The results are presented in the Additional File 1.

### Evaluation using long-reads reference

For the second test with long-read data we used publicly available data from <u>Jin et al. (2022)</u>, corresponding to human fecal samples sequenced by both Illumina HiSeq X Ten platform with 150bp-long reads (SRR10917786, 34.2 Gbp) and PacBio RS II (SRR10917776, 8.7 Gbp) (Pacific Biosciences of California, Inc., USA). We performed error-correction of the raw reads using Canu (<u>Koren et al. 2017</u>):

canu -correct -PacBio-raw SRR10917776.fastq -p PacBio_corrected -d corrected_reads genomeSize=100m

To avoid creating erroneous contigs we decided to not assemble the long-read data but instead rely on the long error-corrected reads as they most likely represent the ground truth. First, we annotated the reads using BLASTN against the ResFinder database (2021) (<u>Florensa et al. 2022</u>). Only reads containing full ARG sequences with at least 98% identity were retained. These reads were further clustered with cd-hit (<u>Fu et al. 2012</u>) (95% identity) to create a less redundant reference for further comparison to the short-read contigs. We assembled the corresponding short reads with the same set of tools as mentioned in the previous section. The resulting contigs were mapped using BLASTN to the PacBio reads reference to assess accuracy.

To estimate how the differences in ability to assemble ARGs by different tools affect the downstream results, we performed ARG quantification. First, the Illumina reads were mapped to each assembly using bowtie2 v2.3.5.1 (<u>Langmead and Salzberg 2012</u>) and the coverage was estimated for all full and truncated ARG hits (minimum 95% identity and 80% coverage) on the assembled contigs using the FARAO (<u>Hammarén et al. 2016</u>)

*estimate_coverage* function with -c 0 flag estimating coverage per all bases across the feature:

estimate_coverage -i assembly_coverage_db -a get_annot_output_filtered -c 0

All plots were created in R using ggplot2 (Villanueva and Chen 2019).

### Assessment

To assess the performance of the tools, we identified contigs produced from the short-reads containing ARG sequences and estimated the number of fully assembled (100% coverage, 98% identity), truncated (minimum 60 bp length, 98% identity and no flanking regions on either of the sides or both sides) and misassembled/partial ARGs (minimum 60 bp, 98% identity, and embedded in incorrect flanking sequences). The 60 bp threshold was chosen because it corresponds to at least 20 amino acids, which should be sufficient to identify a protein. Furthermore, we investigated the genomic contexts of the fully assembled ARGs and whether they fully matched the original context by inspecting alignments to the original plasmids.

## Results

### Assembly quality

We compared the assembly performance of Velvet, Ray, MEGAHIT, metaSPAdes and Trinity (Table 2, Figure S1). Trinity outperformed all the tools for the total assembled length and the number of reconstructed contigs at all coverages. In contrast, metaSPAdes assembled the longest contigs except for at the 10x coverage, where the length of the longest contig was comparable to the one assembled by Ray. metaSPAdes also had the highest and most consistent mapping rate across different assemblies (Table 3).

**Table 2.** Assembly summary statistics.

| Coverage | Assembly |  |  |  |  |  |
| --- | --- | --- | --- | --- | --- | --- |
| 0.5x | Velvet | Ray | SPAdes | MEGAHIT | metaSPAdes | Trinity |
| # contigs | 32,218 | 17,350 | 39,226 | 51,677 | 41,734 | 59,194 |
| Largest contig | 25,401 | 474,180 | 172,683 | 50,052 | 688,925 | 126,557 |
| Total length | 36,029,448 | 37,341,447 | 76,326,026 | 66,984,094 | 70,838,814 | 94,470,748 |
| N50 | 1,231 | 7,066 | 4,253 | 1,591 | 2,838 | 2,382 |
| 1x | Velvet | Ray | SPAdes | MEGAHIT | metaSPAdes | Trinity |
| # contigs | 22,246 | 17,421 | 45,074 | 37,579 | 41,864 | 59,258 |
| Largest contig | 52,173 | 442,619 | 184,495 | 62,496 | 688,925 | 120,871 |
| Total length | 26,584,994 | 37,998,115 | 76,946,301 | 47,137,002 | 71,305,149 | 94,588,910 |
| N50 | 1,318 | 7,254 | 2,893 | 1,492 | 2,865 | 2,388 |
| 5x | Velvet | Ray | SPAdes | MEGAHIT | metaSPAdes | Trinity |
| # contigs | 22,686 | 17,364 | 42,987 | 51,749 | 41,505 | 59,108 |
| Largest contig | 56,951 | 303,232 | 172,683 | 66,255 | 542,864 | 120,871 |
| Total length | 27,444,611 | 37,964,392 | 77,214,295 | 67,927,289 | 72,257,839 | 96,105,718 |
| N50 | 1,355 | 6,760 | 3,262 | 1,629 | 3,063 | 2,470 |
| 10x | Velvet | Ray | SPAdes | MEGAHIT | metaSPAdes | Trinity |
| # contigs | 32,332 | 17,048 | 41,502 | 51,459 | 41,392 | 58,988 |
| Largest contig | 56,951 | 467,217 | 172,683 | 91,584 | 464,577 | 168,064 |
| Total length | 37,378,587 | 41,056,751 | 76,752,084 | 67,809,374 | 72,108,940 | 96,703,227 |
| N50 | 1,295 | 11,883 | 3,485 | 1,638 | 3,080 | 2,520 |
| Real data scenario | Velvet | Ray | SPAdes | MEGAHIT | metaSPAdes | Trinity |
| # contigs | 110,673 | 48,047 | 85,453 | 105,763 | 92,846 | 156,257 |
| Largest contig | 47,990 | 582,509 | 556,300 | 295,746 | 400,849 | 78,113 |
| Total length | 151,747,721 | 145,421,628 | 242,507,147 | 232,009,297 | 239,314,901 | 284,725,193 |
| N50 | 1,752 | 15,515 | 12,135 | 5,044 | 8,114 | 3,046 |

**Table 3.**
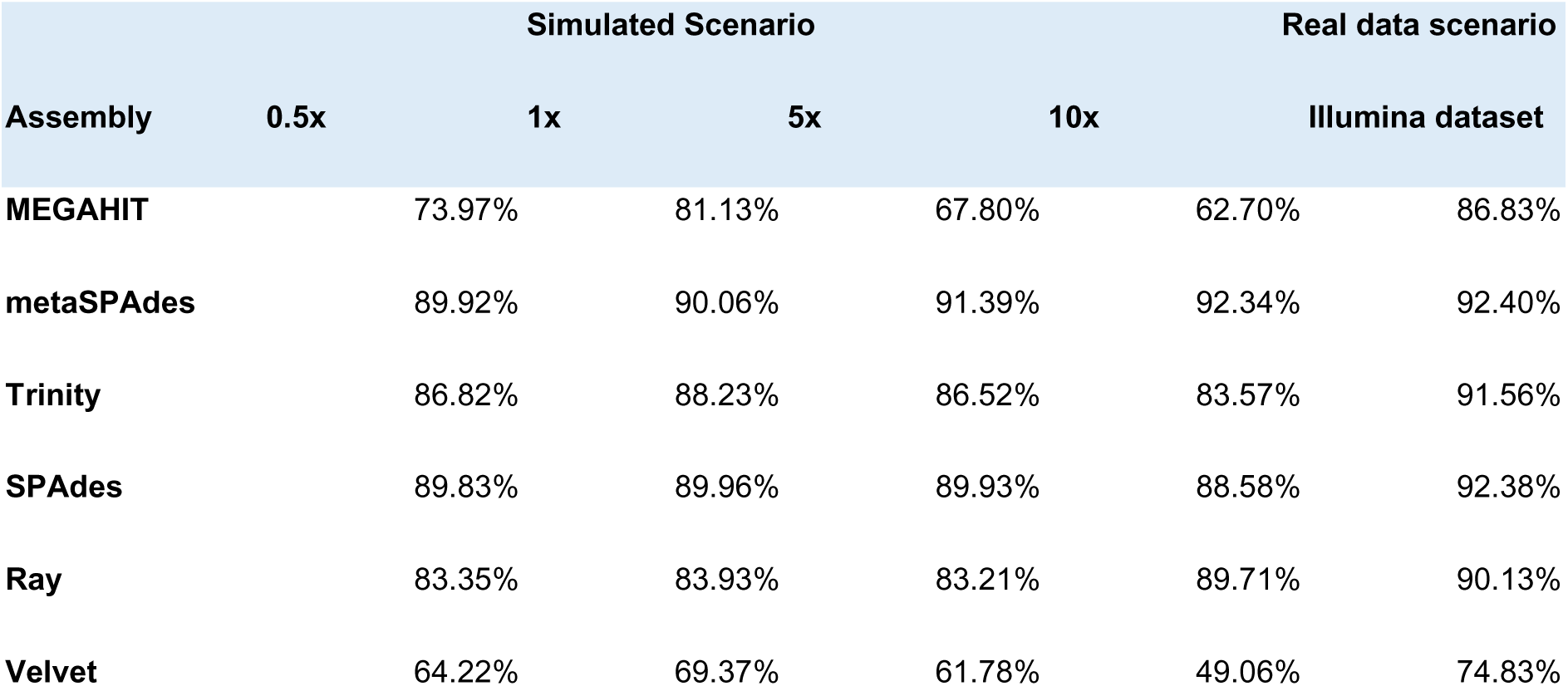
Mapping rate for the simulated scenario and the real dataset.

### Complex short-read data yield incomplete assembly of ARGs and their contexts

To investigate which assemblers managed to reconstruct ARGs from short-read data, we first looked at the recovery of both full length and truncated ARG sequences (Figure 2). For the simulated test data, the knowledge of exactly which ARGs were present on the original plasmids allowed us to precisely determine how many of those were correctly recovered by each assembler (Figure 2A; “presence/absence”). For this analysis, we were interested in whether an ARG was recovered, even if it was assembled in the wrong context, because for some applications it is sufficient to just obtain the individual gene sequence correctly, regardless of context.

**Figure 2.**
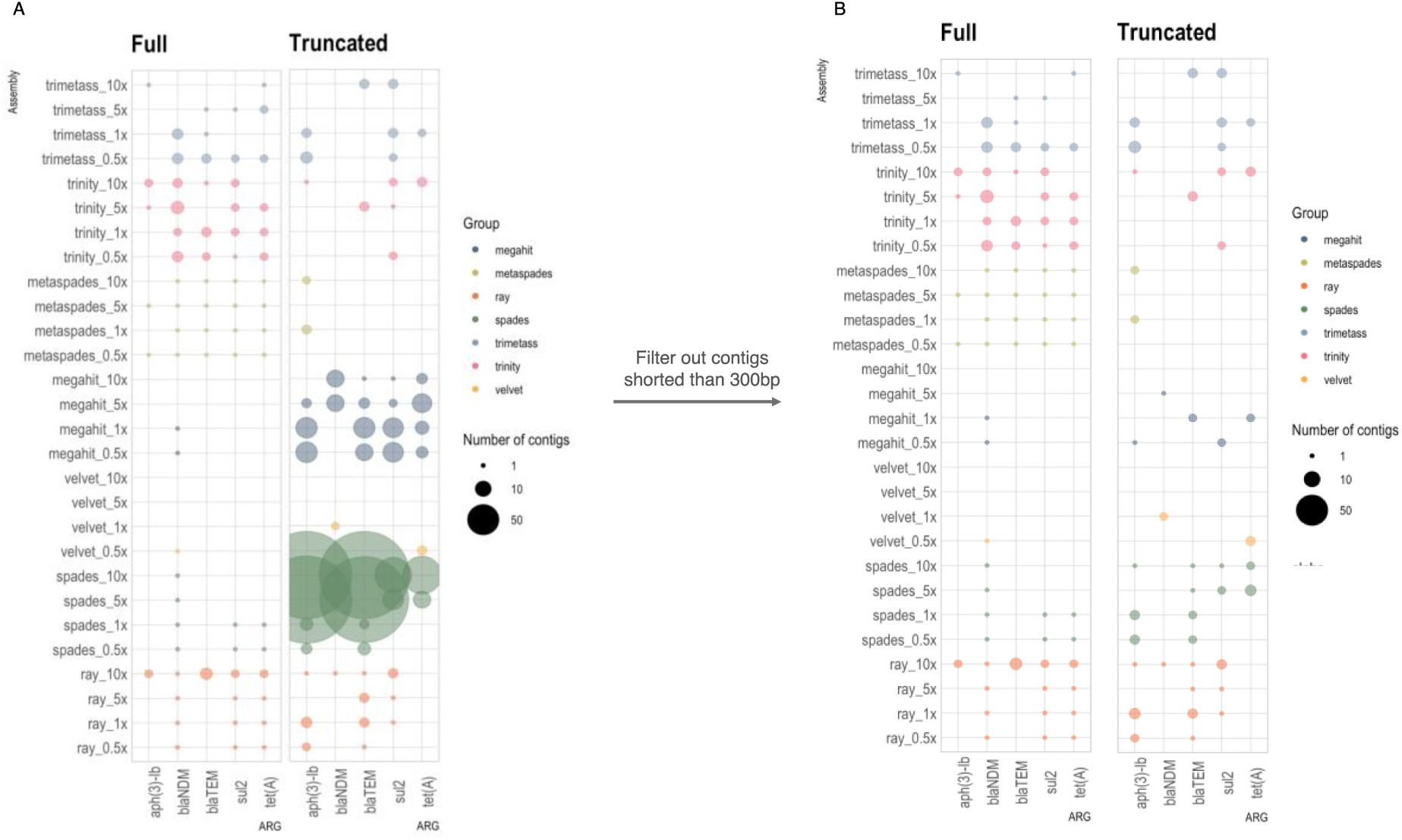
ARGs recovery by each tool: A) presence/absence of ARGs on the contigs assembled by different tools; B) filtering using a length cut-off of 300bp was applied to the results. “Full” denotes contigs containing full length and correctly assembled ARGs while “Truncated” comprises contigs containing partial ARG sequence (minimum 60bp and 98% identity and no flanking regions on either of the sides or both sides).

The results showed that MEGAHIT, metaSPAdes and Trinity managed to capture almost all ARGs at all coverages. However, the MEGAHIT contigs containing ARGs were on average only 284 bp long, resulting in predominantly truncated ARG sequences (Figure 2 and 3). In contrast, Trinity performed consistently better at all coverages, with more than 50% of contigs containing the full length ARG sequences (Figure 3). Among the genome assemblers, Ray had the best performance in terms of reconstructing full ARGs, while Velvet reconstructed only one full ARG out of 3724 assembled ARG-containing contigs, with the rest containing misassembled ARGs. SPAdes struggled to assemble ARGs at higher 314 coverages, producing truncated contigs.

**Figure 3.**
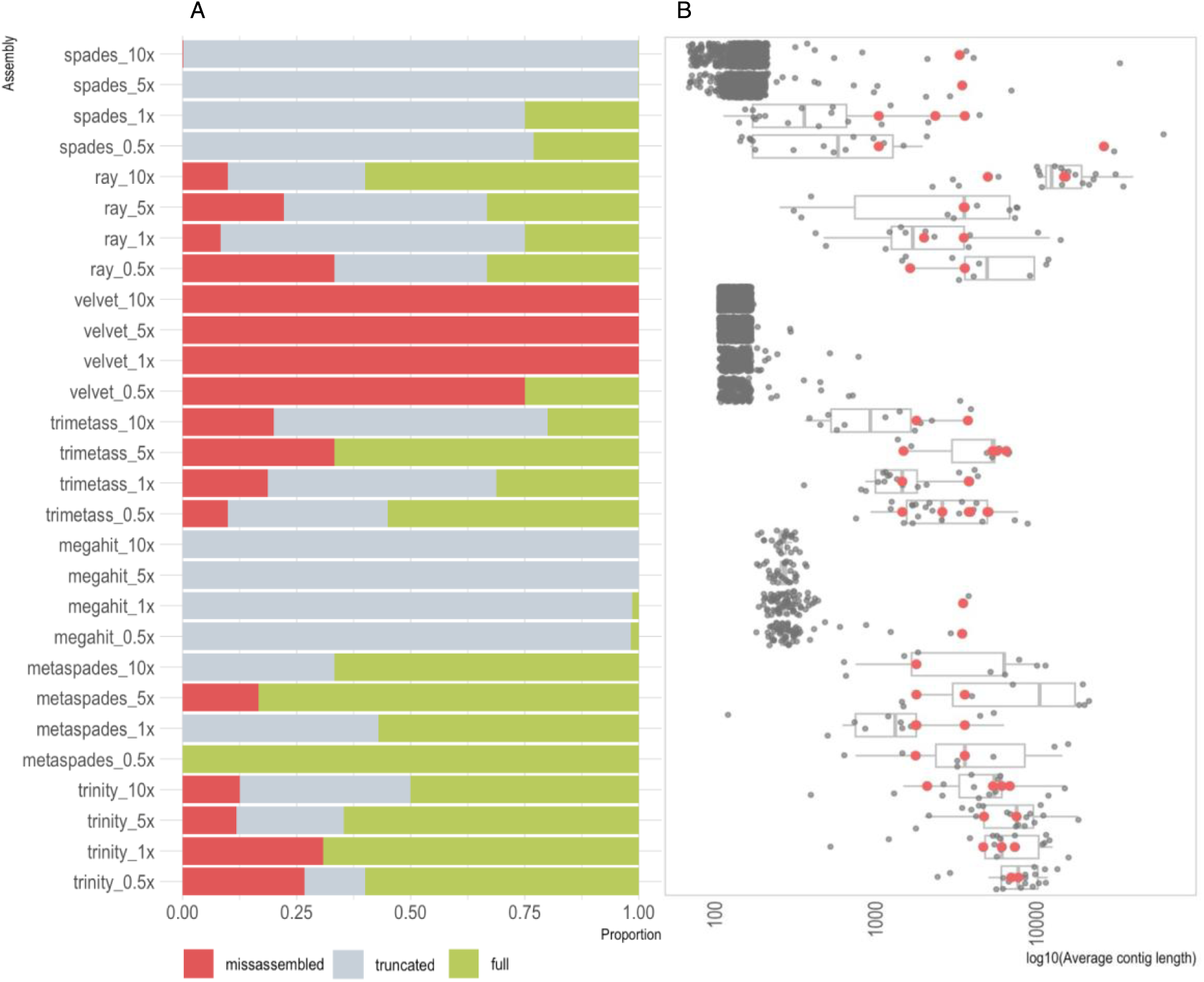
Assembler performance at different coverages: A) proportion of full, truncated and misassembled/partial ARG sequences. Note that the retrieved ARGs are not necessarily associated with the correct context.); B) length distribution of contigs with ARG hits (contigs with correct genomic context, only containing full ARGs, are marked with red dots).

Furthermore, we investigated the number and length of correctly assembled contexts for the different assemblers. Figure 3B shows that Trinity performed better in comparison to the other tools, reconstructing on average longer correct contigs with quite consistent performance across the coverages. Notably, the performance of Ray was rather similar to that of metaSPAdes. In some cases, Ray produced even longer correct contigs despite being a genomic assembler not optimized for complex metagenomic samples. In contrast, MEGAHIT produced only two correct contigs at lower coverages. What is obvious and rather surprising is that from a total number of 5 different genomic contexts per resistance gene present in the sample on average only three original contexts (12%) were correctly captured by any of the assembler. These contexts corresponded to plasmids of different sizes suggesting that the total length of plasmids did not determine assembly success, but rather features surrounding the particular genes (Figure S2).

We also used SPAdes contigs as seeds to extend them using TriMetAss. The results revealed that in general TriMetAss output a few more correctly assembled contigs containing full ARGs. Importantly, these contigs were on average 2000 bp longer than initial SPAdes contigs. As a drawback, TriMetAss also produced more misassembled contigs in comparison to SPAdes output (Figure 3A).

### ARG regions are particularly prone to poor assembly quality

All assemblers reconstructed large contigs spanning in some cases half of the plasmid sequence (as shown on the example of AP023079 plasmid; Figure 4), but broke exactly at the beginning of ARG sequence. The complexity of assembly graphs also increased with more coverage, but for some tools, such as SPAdes and metaSPAdes, additional coverage helped to resolve ambiguous branching and reconstruct longer contigs, while for MEGAHIT increased coverage resulted in profound fragmentation. The assembly graphs also made it clear that Trinity, as a transcriptomic assembler, utilizes a different approach in comparison to the other tools, resulting in very characteristic assembly graph patterns.

**Figure 4.**
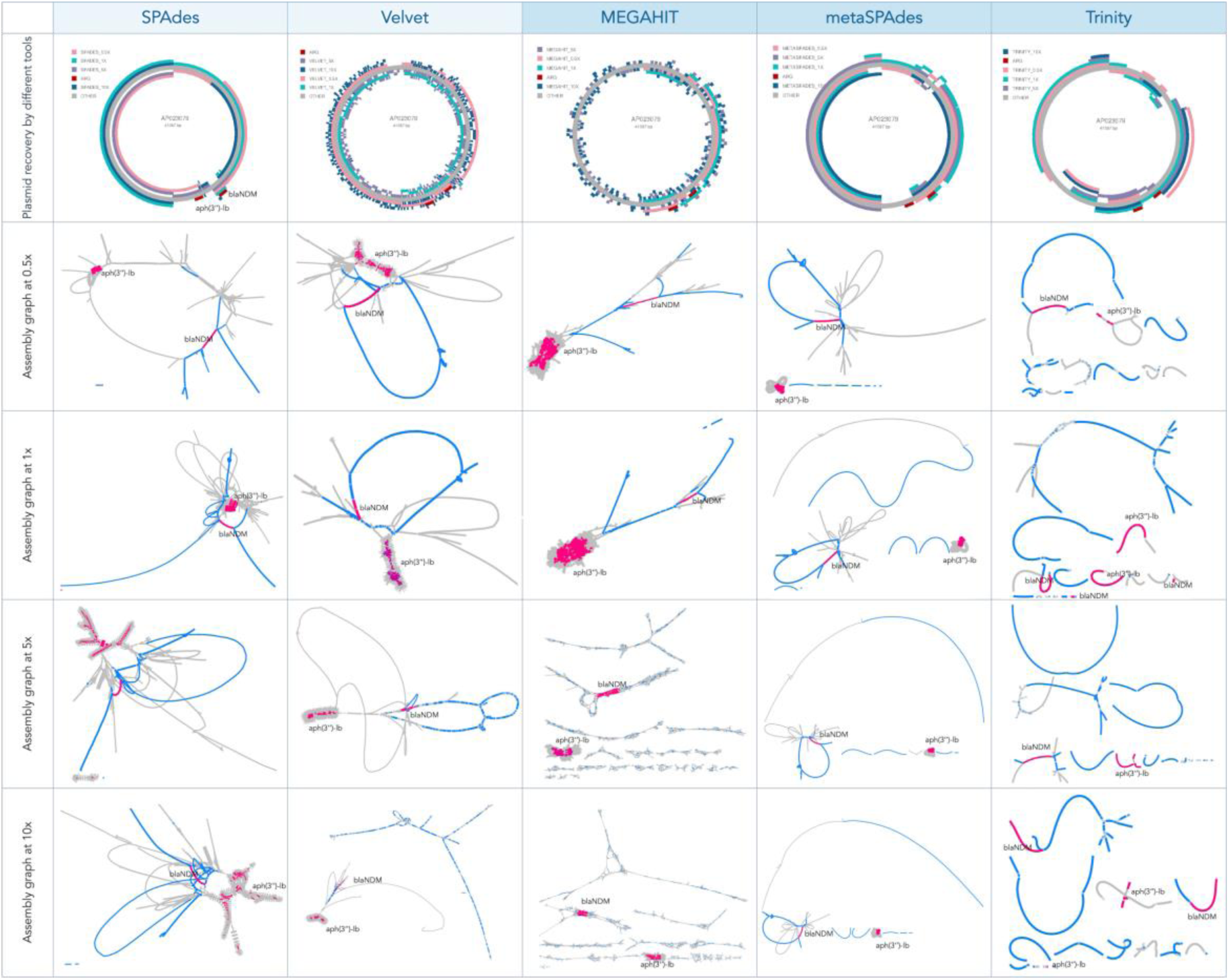
Visual representation of assembly results on the example of one of the plasmids AP023079 containing two ARGs *blaNDM* and *aph(3’’)*: A) visual representation using FARAO: light gray represents the backbone plasmid and the other colors represent correctly assembled contigs from different assemblies (0.5x in pink, 1x in teal, 5x in purple and 10x in blue) and ARGs are in red; B) visual representation of the corresponding assembly graphs using Bandage: the figures represent only part of the whole assembly graph corresponding to the AP023079 plasmid sequence, where blue lines correspond to BLAST hits of the assembled contigs to the plasmid and pink lines to the ARG regions.

To investigate what consequences this fragmentation has on the ARG quantification, we mapped reads back to the corresponding assemblies to estimate gene abundances. In parallel, we quantified ARGs by mapping reads directly to the ResFinder database (Figure 5A) and to the same database but clustered by 90% identity to reduce variant redundancy (Figure 5B).

**Figure 5.**
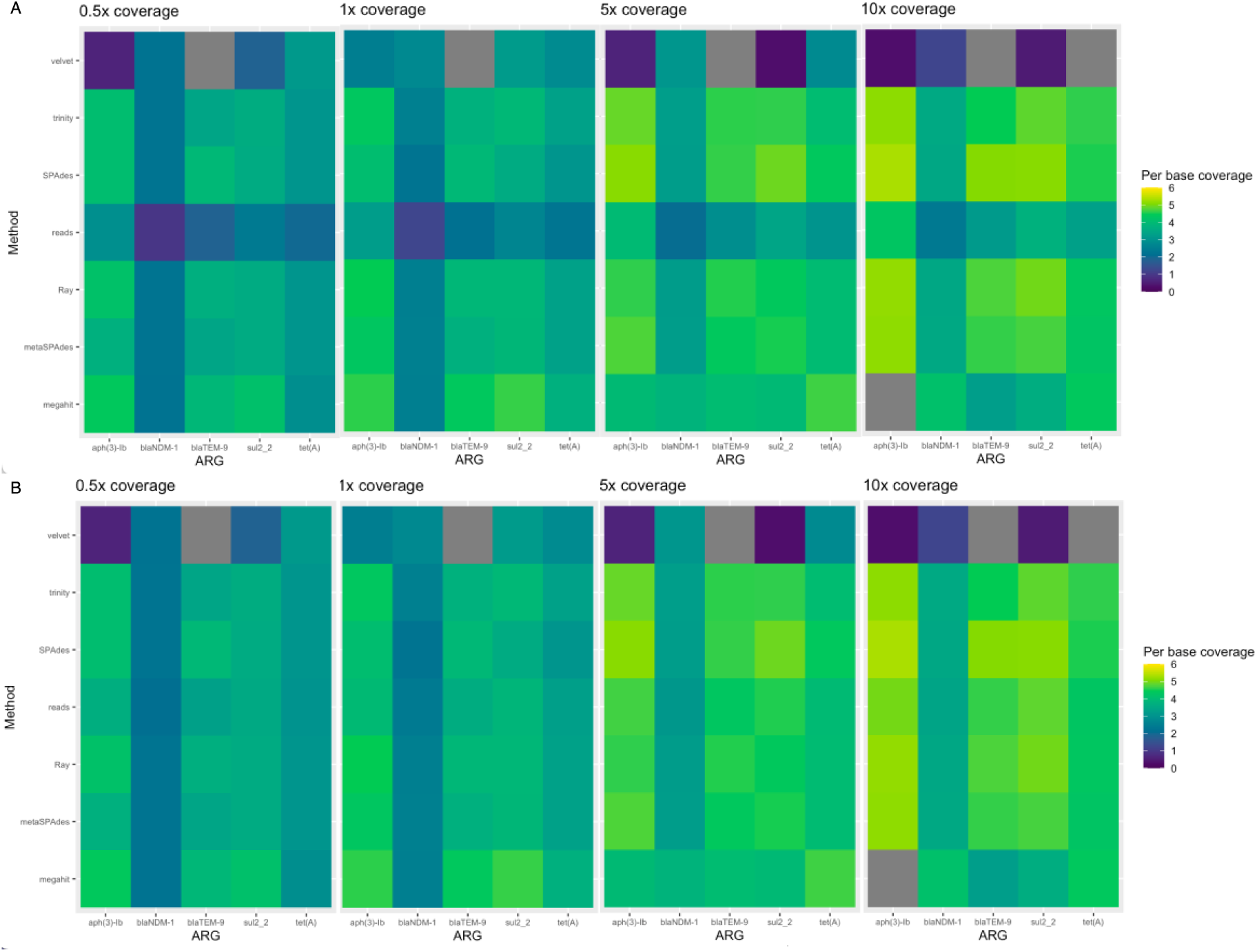
ARG quantification using either assembled contigs as a reference or by directly mapping short reads to the ResFinder database: per base total coverage calculated using FARAO from aligning reads to the contigs, and using direct ARG quantification by mapping reads to ResFinder database (A) and ResFinder clustered to 90% identity (B).

### Long-read data confirmed results of simulated metagenomes

To validate the findings from the simulated metagenome data, we performed a second test with a real dataset. For this, we used PacBio reads containing full ARGs as a reference for the contigs assembled from a corresponding short-read data set derived from the same samples (see Materials and Methods for details). In this dataset, we annotated ARGs on the PacBio reads, which resulted in 18 unique ARGs (98% identity and 100% coverage) found on 125 PacBio reads (32 reads carried more than one ARG). This set of PacBio reads was used as a reference to evaluate contigs containing ARGs assembled from short-read data.

After assembling the short-reads data, we compared these contigs to the PacBio reference reads to assess the correctness of genomic context. In this comparison, Trinity had the highest number of correct contigs matching the reference PacBio reads, and these contigs were on average longer than the ones reconstructed by the other tools (Figure 6A and Table S1). In contrast, MEGAHIT, metaSPAdes and SPAdes assembled half as many contigs with on average shorter length than Trinity. To check if another approach using TriMetAss could improve the results, we used SPAdes contigs containing full and truncated ARGs as a seed for iterative re-assembling. However, the results revealed that this approach did not considerably improve the length of the contigs containing ARGs.

**Figure 6.**
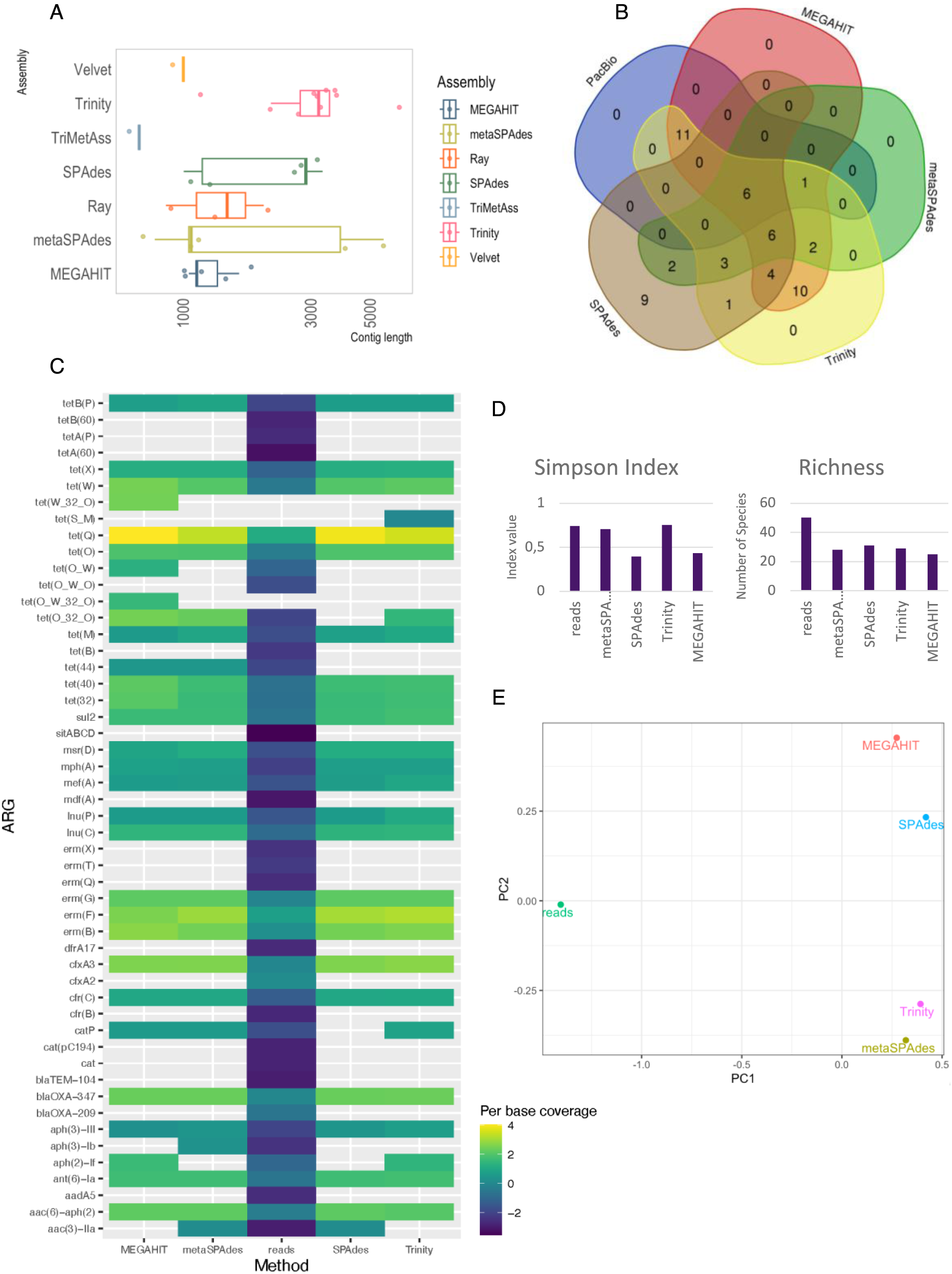
Results from real case scenario: A) Length distribution of contigs assembled from short reads matching the PacBio reference reads; B) Number of unique ARGs identified on assembled contigs and PacBio reference reads (Ray, Velvet and TriMetAss are not shown); C) ARG quantification using assembly reference and direct mapping of short reads: FARAO results for assemblies and direct ARG quantification by mapping reads to the ResFinder database, D) Simpson diversity and Richness (unique ARGs identified); E) PCoA based on Bray-Curtis distances between quantification methods.

### Long-read sequencing does not reliably detect all ARGs

All together, 55 unique ARGs were identified from all the assembled contigs by all the short-read tools and PacBio reads, with only 6 ARGs common between all of them (Figure 6B). Interestingly, Trinity, SPAdes and metaSPAdes recovered several additional ARGs (full or truncated) not present on PacBio reads. Importantly, annotation of short reads alone resulted in 85 matches (80% coverage and 98% identity).

To assess how this discrepancy in ARG identification would affect the results of ARG quantification, we separately mapped short-reads to assembled contigs, as well as to the ResFinder database to quantify ARG abundances directly from the reads. This provides a direct comparison between the two prevailing approaches to quantify ARGs in metagenomic data (Bengtsson-Palme et al. 2017). This analysis revealed that for some ARGs, results are very similar between approaches, such as for several tetracycline genes (e.g., *tetQ* and *tetW*), as well as the erythromycin resistance gene *ermB* and aminoglycoside resistance gene *aph(3’)-III* (Figure 6C). However, several ARGs were detected only by mapping reads directly to the ResFinder database and were missing completely from the assembly-based approach (e.g., *tetB*, *blaTEM-104* and *ermT*).

## Discussion

### The repertoire of ARGs recovered from the same dataset differs depending on the tool used

It is commonly assumed that metagenomic high-throughput sequencing allows an *unbiased* cataloging of ARGs at the whole microbiome scale (<u>Su et al. 2017</u>; <u>Lee et al. 2021</u>) in comparison to qPCR and culture-based methods. However, it is rarely considered that downstream data processing can have major impacts on the results reported. There have been several initiatives to benchmark metagenomics software. <u>Wang et al. (2019)</u> performed evaluation of metagenomic assemblers using real metagenomic datasets spiked with reads from known genomes focusing on completeness and accuracy of reconstructed genomes. A few studies have looked into the benefits of using long-reads for improving metagenomic assemblies (<u>Bertrand et al. 2019</u>; <u>Latorre-Pérez et al. 2020</u>; <u>Xie et al. 2020</u>). Within the framework of the CAMI challenge, <u>Sczyrba et al. (2017)</u> and <u>Meyer et al. (2022)</u> evaluated assembly performance, but largely for the purpose of taxonomic profiling. However, only a few studies have looked into the implications of assembler choice for the inference of gene contexts and gene abundances from metagenomic assemblies. <u>Brown et al. (2021)</u> used resistome risk score based on co-occurrences of ARGs, MGEs and pathogen gene markers on the same contig to evaluate convergence of biological output produced by different assemblers. Another paper by <u>Galata et al. (2021)</u> showed that the choices of assembly software as well as sample complexity have considerable impact on prediction of genes and proteins. A recent study by <u>Yorki et al. (2023)</u> focused on assessment of short-, long- and hybrid-approaches to recover the genome of clinically relevant low-abundant *E. coli* and their ARG content from metagenomic samples. Unfortunately, the results of these studies are often contradictory, most probably due to the different approaches and features of the underlying test data. Most importantly, despite being mentioned by several studies, the impact of assembler choice on the biological interpretability has not been well explored.

In the current study, we used both simulated and real metagenomics data to assess the impact of assembler choice on the identification and quantification of ARGs, as well as the ability to correctly reconstruct the genomic contexts surrounding these ARGs. Metagenomic assemblers are optimized to deal with sequence data from samples containing multiple species in different abundances, and therefore their performance was of primary interest. In the simulated scenario, metaSPAdes considerably outperformed MEGAHIT in terms of number of contigs containing full ARG sequences, with MEGAHIT predominantly producing short contigs (on average 284 bp long) with truncated ARGs. This could have severe consequences on the final results since many metagenomic studies utilizing the assembly approach employ a filtering step to remove short, potentially erroneous, contigs containing little useful information. The filtering cut-off can vary from 300 bp up to 2 kb (<u>Dang et al. 2020</u>; <u>Chen et al. 2022</u>; <u>Yi et al. 2022</u>; <u>Ke et al. 2023</u>). Even if we were to apply the most allowing cut-off of 300 bp to the MEGAHIT results, 99% of contigs containing ARGs would be filtered out, resulting in a considerable underestimation of the resistome in the sample. This suggests that the choice of assembler as well as pre-processing and post-processing steps can considerably affect the end result.

This problem became even more profound when we performed ARG identification in a real dataset. For a real dataset, there is no way to know the true complete repertoire of ARGs. Therefore, we used PacBio reads containing full ARGs as a reference for the contigs assembled from corresponding short-read data. All together, 55 unique ARGs were identified from all the assembled contigs across all the short-read tools and the PacBio reads (Figure 6B). The number of unique ARGs captured by the different assemblers varied greatly, from 44 identified by Trinity to 20 captured by metaSPAdes. Similarly to the assemblies from the simulated data, 52% of the MEGAHIT contigs containing ARGs would have been filtered out using a 300 bp length cut-off, showing that this undesired effect is not simply a matter of our methodological choices for simulating data. Interestingly, Trinity, SPAdes and metaSPAdes recovered several additional ARGs not present on the PacBio reads. In this particular case, the short read dataset was sequenced three times deeper than the PacBio dataset, suggesting that the long read dataset did not have sufficient depth to pick up all the ARGs. That said, the two approaches would probably perform similarly well at comparable sequencing depth, but the costs of long read sequencing would – at present – be considerably higher.

Worryingly, only six ARGs were commonly identified by all tools, including two aminoglycoside, two tetracycline and two erythromycin resistance genes. Consequently, some genes were missing from the output by all short-read assemblers tested, including the clinically relevant beta-lactamase gene *blaOXA*, which was identified on PacBio reads and therefore most certainly present in the sample. The most probable explanation for this is that those were rare genes that did not get sufficient coverage to be assembled by the short-read sequencing effort and therefore are missing from the resulting assembly. An alternative approach to ARG quantification in metagenomes, circumventing assembly, is identification of ARGs by mapping the reads directly to one of the available ARG databases (<u>Bengtsson-Palme et al. 2017</u>). We annotated reads by mapping reads to the ResFinder database, which resulted in identification of 85 unique ARGs. Perhaps not surprisingly, this number by far surpassed the total number of ARGs identified by mapping reads to the assembled contigs, as well as on PacBio reads alone. Reads are typically much shorter than contigs and might map spuriously to several different targets causing false positives. At the same time, this approach does not require coverage of the entire ARG in order to detect it, which may be crucial for the detection of rare ARGs. As many clinically relevant ARGs to last resort antibiotics are typically rare in most microbiomes, the increase of detection ability is highly important for e.g. monitoring of high-risk ARGs (<u>Abramova et al. 2023</u>; <u>Bengtsson-Palme et al. 2023</u>). This finding also highlights the importance of not basing gene catalogs only on assemblies from the metagenomes under study, but also including relevant gene or genome repositories into the catalogs used for annotation and read mapping.

Depending on which tool and cut-off are used for the data analysis, the end results can be drastically different. It is important to mention though that the number of correctly assembled full-length ARGs on its own is not always a good measure of assembler performance, if the rest of the output contigs contain misassembled sequences. In most real-world scenarios, it would not be possible to determine which contigs were correctly and incorrectly assembled, underscoring the importance of assembly tools achieving a good ability to stitch reads together while still maintaining strict precision in terms of obtaining the correct assembled contexts.

### Correctly assembled short contigs often lack context around ARGs

Obtaining a correctly assembled full or even truncated ARG might be enough for certain applications, for example when estimating the ARG diversity in a sample. However, for the purpose of host taxonomic inference or mobility assessment of a given ARG, it is necessary to look into the genomic context around it. After assembling short-reads data we compared the resulting contigs to the original plasmid sequences for a simulated data set or to PacBio reference reads for the real data scenario, allowing us to assess the correctness of the assembled genomic contexts. In this comparison, Trinity had the highest number of correct contigs matching the reference in both cases, and the Trinity contigs were on average longer than the ones reconstructed by the other tools (Figure 3B and Table S1). Trinity is a transcriptome assembler, specifically designed to assemble transcript variants resulting from alternative splicing or gene duplication (<u>Grabherr et al. 2011</u>). Instead of trying to reconstruct the full graph, it starts with assembling disjoint transcription loci which are further converted into de Brujin graphs and pruned based on read support. The difference to the other assembler approaches is visible in the graph representation (Figure 4), where Trinity contigs are represented by nodes of more even coverage and less complexity in comparison to graphs resulting from metaSPAdes and MEGAHIT assemblies. The original MEGAHIT publication (<u>Li et al. 2015</u>) showed that its performance becomes better with increased coverage (from 10x to 100x) in terms of N50 value, largest alignment length and number of misassemblies. However, we observed that in our simulated data scenario MEGAHIT performed best at lower coverages (0.5x and 1x; Figure 2 and 3), showing extensive fragmentation at the higher coverages as revealed by the highly branching graph (Figure 4). However, in the real data scenario MEGAHIT and metaSPAdes showed very similar performance in terms of number of correct contigs and their length. This is somewhat reflective of our simulated approach representing a very complex, but yet realistic, case in terms of the number of different resistance plasmids present in the simulated data. That said, due to the rather short length of the MEGAHIT and metaSPAdes contigs they match to several different PacBio reads (different genomic contexts) implying that their length is not sufficient to unambiguously decipher the taxonomic origin of the ARGs they carry. Taking into account that the average ARGs is longer than 500 bp, these contigs most probably also lack any information about co-located ARGs or MGEs, at least with any degree of certainty.

### Quantification of ARGs is heavily dependent on correct assemblies

As has been discussed above, the repertoire of ARGs detected in the assemblies varied greatly between different assembly tools in both scenarios. To investigate what consequences this has on the ARG quantification, we mapped reads back to the corresponding assemblies to estimate gene abundances. In parallel, we quantified ARGs by mapping reads directly to the ResFinder database. For the simulated scenario, we have knowledge about exactly which ARG sequences should be present in the dataset and therefore can directly compare abundances for these particular genes estimated by two different approaches. It was surprising to observe that quantification by mapping reads back to the ResFinder database revealed in general lower abundance levels than when calculating abundance by mapping to the assembled contigs (Figure 5A). The ResFinder database, as well as the other AMR gene catalogs, contains two hierarchical levels of nomenclature: gene family such as *blaNDM* or *tet(M)* and their associated allelic variants (e.g. *blaNDM-1*, *tet(M)-6*). We hypothesized that the observed results is a consequence of variant “spill-over” effect when the lengths of the reads were insufficient to differentiate between the variants.

A possible solution often used to reduce the impact of this variant “spill-over” effect is to cluster all the similar variants and retain only a single representative sequence. We did this in an additional test using the ResFinder database clustered to 90% identity (Figure 5B). This approach allowed us to estimate the abundance for a family of closely related ARG variants instead of a particular variant. Despite the somewhat lower resolution, using a clustered database yielded abundance estimates for all the spiked-in genes comparable to those estimated based on the mapping to assembled contigs.

In the real-case scenario, the results revealed that several ARGs were quantified only by mapping short reads directly to the ResFinder database and were missing completely from the assembly-based approach (Figure 6C). Some of those are clinically relevant genes such as *tetA*, *tetB*, *blaTEM* and *ermT*, and therefore, it is crucial to understand if their presence/absence is an artifact. In the case of *blaTEM*, it was only identified by mapping the short reads to the ResFinder database, and it was missing from both PacBio reads as well as assembled contigs. A closer inspection of the original Illumina dataset showed that it is a low-abundant gene that did not have enough coverage to be assembled by short-read assemblers and was not picked-up by long-read sequencing.

All together, these results show that none of the approaches give a comprehensive picture of ARG diversity and abundance. Assembled contigs provide a good resolution in terms of identification and quantification of specific ARG variants as well as their genomic context, but at the same time this approach misses rare ARGs due to insufficient coverage. In contrast, direct mapping of short reads spuriously aligns them to different ARG variants, leading to an overestimation of the resistome diversity in a sample. There are several studies suggesting the importance of knowing the genomic context of ARG variants for determining their transmission potential, co-resistance patterns and how well they would respond to different interventions (<u>Zhang et al. 2021</u>; <u>Munk et al. 2022</u>). Therefore, using read-based quantification alone to determine ARG abundance in a sample can result in a misleading interpretation regarding which particular variant is present and abundant in a sample. This highlights the importance of using a combination of approaches to obtain an unbiased picture for ARG diversity and abundance in a metagenomic sample and to exercise caution when interpreting individual ARG results from metagenomic data.

### Certain ARG contexts are particularly hard to assemble correctly

Interestingly, not all of the ARGs were equally easy to assemble (Figure 2), with *aph(3’’)-lb* being the most difficult gene to fully assemble among the ones spiked-in. This is a good example of what happens during assembly of regions which are present in multiple genomic contexts with differential coverage in the same sample. On the original plasmids, the aminoglycoside resistance gene *aph(3’’)-lb* was surrounded by insertion sequences, several other ARGs and recombinases, all contributing to making it difficult to assemble the region around the gene correctly. Not surprisingly, the assembly graphs showed that this problem becomes more pronounced with increasing coverage (Figure 4, the brush-like structures representing *aph(3’’)-lb*). In a nutshell, this problem is analogous to the recovery of 16S rRNA genes from metagenomic samples. The 16S rRNA is a gene consisting of a patchwork of hypervariable and universally conserved regions, resulting in highly complex branched assembly structures (<u>Vollmers et al. 2017</u>).

A factor further complicating metagenomic assembly is that microbiomes are typically characterized by different abundance levels of various species and as a result DNA sequencing yields a highly non-uniform distribution of read coverages across different genomes. In addition, read coverages for most species are much lower than in a typical cultivated single-species sample. All together, these features of metagenomic data cause standard genome assembly procedures to produce fragmented and error-prone assemblies, as can be seen in the examples of Velvet, Ray and SPAdes.

## Conclusions

Overall, there is a need for better assembly software to deal with ARGs in multiple contexts, as the results of this study show that none of the current tools are capable of dealing with samples of high complexity. Currently available metagenomics assembly tools metaSPAdes and MEGAHIT are able to identify a variety of ARGs, but fail to fully recover the diversity of genomic contexts present in a sample. The transcriptomic assembler Trinity, despite being designed for a different purpose, is an interesting alternative as it showed better performance in reconstructing longer and fewer contigs matching unique genomic contexts, which can be beneficial for deciphering the taxonomic origin of ARGs. Therefore, for situations where a complex metagenome can be expected, we would recommend using Trinity. However, often the available computational resources will not allow this, as Trinity is a computationally very demanding software. As a second option, we suggest metaSPAdes, which also requires a lot of resources, and therefore is feasible only for smaller datasets, unless substantial computational resources are available. MEGAHIT showed quite poor performance in a complex case scenario, producing very short contigs, but its performance was comparable to that of metaSPAdes in the real data scenario. We would suggest MEGAHIT as an option for low complexity samples as it is also much more CPU- and memory-friendly than all other approaches. In addition, MEGAHIT might sometimes be the only feasible option for producing assemblies from very large datasets. This is not an ideal situation from a point of view of assigning contexts to ARGs and highlights the necessity of developing new approaches to short-read assembly.

Finally, we have made one very important observation: our results show that using a length filtering threshold for the assembled contigs can contribute to a dramatic loss of ARG-containing contigs. This is due to that ARGs seem to be over-represented among challenging genomic contexts for assembly, and for that reason these regions are particularly prone not to be properly assembled, resulting in short and fragmented contigs. This can lead to drastic underestimation of the resistome diversity and abundance in a sample. We suggest, therefore, to annotate ARGs on contigs *before* filtering on the length, to have an idea of what is being filtered out. When it comes to ARG abundance quantification, direct mapping of ARGs to a database rather than an assembly results in better detection ability, but risks increasing false positive detections. One alternative approach is to cluster the reference database to reduce the number of ARG variants. This will lead to a lower resolution at the ARG variant level, but will on the other hand reduce the risks for biasing the picture of ARG prevalence. Another way could be to assign a threshold for the minimal number of reads mapped, distributed across a reference ARG, to make sure that there is enough support for that gene being present in a sample.

In conclusion, as researchers we should not blindly trust the output of our bioinformatics tools. If tools corroborate each other, one can put more trust into their output. If not, one should exercise a lot of caution when interpreting data, especially on genetic contexts in potentially complicated regions. Long read sequencing may eventually solve these problems, but we are not there yet, partially because of the excessive costs of deep long-read sequencing. In the meantime, new more accurate methods are needed to resolve the contexts around ARGs in order to determine where they belong taxonomically and their potential for mobility.

## List of abbreviations

ARGs: Antibiotic resistance genes
AMR: Antimicrobial resistance
RNA: Ribonucleic acid
CAMI: Critical Assessment of Metagenome Interpretation
MGEs: Mobile genetic elements
SRA: Sequence Read Archive
bp: base pairs
CARD: Comprehensive Antibiotic Resistance Database
NCBI: National Center for Biotechnology Information
qPCR: Quantitative Polymerase Chain Reaction

## Declarations

### Ethics approval and consent to participate

Not applicable

### Consent for publication

Not applicable

## Availability of data and material

The datasets analysed during the current study are available in the SRA database:

https://www.ncbi.nlm.nih.gov/sra/?term=SRR10917786,

https://www.ncbi.nlm.nih.gov/sra/?term=SRR9654970

## Authors’ contributions

JBP conceptualized, supervised, and acquired funding for the work. All authors contributed to design of the experimental framework. AA produced and analyzed results; wrote the manuscript. All authors contributed to interpretation of the data and revision of the manuscript. All authors read and approved the final manuscript.

## Acknowledgements

The authors would like to acknowledge funding from the Swedish Research Council (VR; grant 2019-00299) under the frame of JPI AMR (EMBARK; JPIAMR2019-109), the Data-Driven Life Science (DDLS) program supported by the Knut and Alice Wallenberg Foundation (KAW 2020.0239), the Swedish Foundation for Strategic Research (FFL21-0174), the Centre for Antibiotic Resistance Research at the University of Gothenburg, the Sahlgrenska Academy at the University of Gothenburg, and the Swedish Cancer and Allergy fund (Cancer-och Allergifonden). We would like to thank Marcus Wenne and Vi Varga for constructive feedback on earlier versions of the manuscript.

## References

Abramova A, Berendonk TU, Bengtsson-Palme J (2023) A global baseline for qPCR-determined antimicrobial resistance gene prevalence across environments. Environment International 178: 108084.

Ayling M, Clark MD, Leggett RM (2019) New approaches for metagenome assembly with short reads. Briefings in Bioinformatics 21: 584–594. doi:10.1093/bib/bbz020.

Bankevich A, Nurk S, Antipov D, Gurevich AA, Dvorkin M, Kulikov AS, Lesin VM, Nikolenko SI, Pham S, Prjibelski AD (2012) SPAdes: a new genome assembly algorithm and its applications to single-cell sequencing. Journal of computational biology 19: 455–477. 10.1089/cmb.2012.0021.

Bengtsson-Palme J, Abramova A, Berendonk TU, Coelho LP, Forslund SK, Gschwind R, Heikinheimo A, Jarquin-Diaz VH, Khan AA, Klümper U (2023) Towards monitoring of antimicrobial resistance in the environment: For what reasons, how to implement it, and what are the data needs? Environment International: 108089. 10.1016/j.envint.2023.108089.

Bengtsson-Palme J, Boulund F, Fick J, Kristiansson E, Larsson DJ (2014) Shotgun metagenomics reveals a wide array of antibiotic resistance genes and mobile elements in a polluted lake in India. Frontiers in microbiology 5: 648. 10.3389/fmicb.2014.00648.

Bengtsson-Palme J, Larsson DJ, Kristiansson E (2017) Using metagenomics to investigate human and environmental resistomes. Journal of Antimicrobial Chemotherapy 72: 2690–2703.

Bertrand D, Shaw J, Kalathiyappan M, Ng AHQ, Kumar MS, Li C, Dvornicic M, Soldo JP, Koh JY, Tong C, Ng OT, Barkham T, Young B, Marimuthu K, Chng KR, Sikic M, Nagarajan N (2019) Hybrid metagenomic assembly enables high-resolution analysis of resistance determinants and mobile elements in human microbiomes. Nature Biotechnology 37: 937–944. doi:10.1038/s41587-019-0191-2.

Boisvert S, Laviolette F, Corbeil J (2010) Ray: simultaneous assembly of reads from a mix of high-throughput sequencing technologies. Journal of computational biology 17: 1519–1533. 10.1089/cmb.2009.0238.

Brown CL, Keenum IM, Dai D, Zhang L, Vikesland PJ, Pruden A (2021) Critical evaluation of short, long, and hybrid assembly for contextual analysis of antibiotic resistance genes in complex environmental metagenomes. Scientific Reports 11: 3753. doi:10.1038/s41598-021-83081-8.

Chen J, Yang Y, Jiang X, Ke Y, He T, Xie S (2022) Metagenomic insights into the profile of antibiotic resistomes in sediments of aquaculture wastewater treatment system. Journal of Environmental Sciences 113: 345–355. 10.1016/j.jes.2021.06.026.

Dang C, Xia Y, Zheng M, Liu T, Liu W, Chen Q, Ni J (2020) Metagenomic insights into the profile of antibiotic resistomes in a large drinking water reservoir. Environment International 136: 105449. 10.1016/j.envint.2019.105449.

Florensa AF, Kaas RS, Clausen PTLC, Aytan-Aktug D, Aarestrup FM (2022) ResFinder–an open online resource for identification of antimicrobial resistance genes in next-generation sequencing data and prediction of phenotypes from genotypes. Microbial Genomics 8:

Fu L, Niu B, Zhu Z, Wu S, Li W (2012) CD-HIT: accelerated for clustering the next-generation sequencing data. Bioinformatics 28: 3150–3152. doi:10.1093/bioinformatics/bts565.

Galata V, Busi SB, Kunath BJ, de Nies L, Calusinska M, Halder R, May P, Wilmes P, Laczny CC (2021) Functional meta-omics provide critical insights into long- and short-read assemblies. Briefings in Bioinformatics 22: doi:10.1093/bib/bbab330.

Gourlé H, Karlsson-Lindsjö O, Hayer J, Bongcam-Rudloff E (2018) Simulating Illumina metagenomic data with InSilicoSeq. Bioinformatics 35: 521–522. doi:10.1093/bioinformatics/bty630.

Grabherr MG, Haas BJ, Yassour M, Levin JZ, Thompson DA, Amit I, Adiconis X, Fan L, Raychowdhury R, Zeng Q, Chen Z, Mauceli E, Hacohen N, Gnirke A, Rhind N, di Palma F, Birren BW, Nusbaum C, Lindblad-Toh K, Friedman N, Regev A (2011) Full-length transcriptome assembly from RNA-Seq data without a reference genome. Nature Biotechnology 29: 644-652. doi:10.1038/nbt.1883.

Haas BJ, Papanicolaou A, Yassour M, Grabherr M, Blood PD, Bowden J, Couger MB, Eccles D, Li B, Lieber M (2013) De novo transcript sequence reconstruction from RNA-seq using the Trinity platform for reference generation and analysis. Nature protocols 8: 1494–1512. 10.1038%2Fnprot.2013.084.

Hammarén R, Pal C, Bengtsson-Palme J (2016) FARAO: the flexible all-round annotation organizer. Bioinformatics 32: 3664–3666. doi:10.1093/bioinformatics/btw499.

Hendriksen RS, Munk P, Njage P, van Bunnik B, McNally L, Lukjancenko O, Röder T, Nieuwenhuijse D, Pedersen SK, Kjeldgaard J, Kaas RS, Clausen PTLC, Vogt JK, Leekitcharoenphon P, van de Schans MGM, Zuidema T, de Roda Husman AM, Rasmussen S, Petersen B, Bego A, Rees C, Cassar S, Coventry K, Collignon P, Allerberger F, Rahube TO, Oliveira G, Ivanov I, Vuthy Y, Sopheak T, Yost CK, Ke C, Zheng H, Baisheng L, Jiao X, Donado-Godoy P, Coulibaly KJ, Jergović M, Hrenovic J, Karpíšková R, Villacis JE, Legesse M, Eguale T, Heikinheimo A, Malania L, Nitsche A, Brinkmann A, Saba CKS, Kocsis B, Solymosi N, Thorsteinsdottir TR, Hatha AM, Alebouyeh M, Morris D, Cormican M, O’Connor L, Moran-Gilad J, Alba P, Battisti A, Shakenova Z, Kiiyukia C, Ng’eno E, Raka L, Avsejenko J, Bērziņš A, Bartkevics V, Penny C, Rajandas H, Parimannan S, Haber MV, Pal P, Jeunen G-J, Gemmell N, Fashae K, Holmstad R, Hasan R, Shakoor S, Rojas MLZ, Wasyl D, Bosevska G, Kochubovski M, Radu C, Gassama A, Radosavljevic V, Wuertz S, Zuniga-Montanez R, Tay MYF, Gavačová D, Pastuchova K, Truska P, Trkov M, Esterhuyse K, Keddy K, Cerdà-Cuéllar M, Pathirage S, Norrgren L, Örn S, Larsson DGJ, Heijden TVd, Kumburu HH, Sanneh B, Bidjada P, Njanpop-Lafourcade B-M, Nikiema-Pessinaba SC, Levent B, Meschke JS, Beck NK, Van CD, Phuc ND, Tran DMN, Kwenda G, Tabo D-a, Wester AL, Cuadros-Orellana S, Amid C, Cochrane G, Sicheritz-Ponten T, Schmitt H, Alvarez JRM, Aidara-Kane A, Pamp SJ, Lund O, Hald T, Woolhouse M, Koopmans MP, Vigre H, Petersen TN, Aarestrup FM, The Global Sewage Surveillance project c (2019) Global monitoring of antimicrobial resistance based on metagenomics analyses of urban sewage. Nature Communications 10: 1124. doi:10.1038/s41467-019-08853-3.

Jin H, You L, Zhao F, Li S, Ma T, Kwok L-Y, Xu H, Sun Z (2022) Hybrid, ultra-deep metagenomic sequencing enables genomic and functional characterization of low-abundance species in the human gut microbiome. Gut Microbes 14: 2021790. 10.1080/19490976.2021.2021790.

Ke Y, Sun W, Jing Z, Zhu Y, Zhao Z, Xie S (2023) Antibiotic resistome alteration along a full-scale drinking water supply system deciphered by metagenome assembly: Regulated by seasonality, mobile gene elements and antibiotic resistant gene hosts. Science of The Total Environment 862: 160887. 10.1016/j.scitotenv.2022.160887.

Koren S, Walenz BP, Berlin K, Miller JR, Bergman NH, Phillippy AM (2017) Canu: scalable and accurate long-read assembly via adaptive k-mer weighting and repeat separation. Genome research 27: 722–736. doi:doi:10.1101/gr.215087.116.

Langmead B, Salzberg SL (2012) Fast gapped-read alignment with Bowtie 2. Nature Methods 9: 357–359. doi:10.1038/nmeth.1923.

Latorre-Pérez A, Villalba-Bermell P, Pascual J, Vilanova C (2020) Assembly methods for nanopore-based metagenomic sequencing: a comparative study. Scientific Reports 10: 13588. doi:10.1038/s41598-020-70491-3.

Lee K, Kim D-W, Cha C-J (2021) Overview of bioinformatic methods for analysis of antibiotic resistome from genome and metagenome data. Journal of Microbiology 59: 270–280. doi:10.1007/s12275-021-0652-4.

Li D, Liu C-M, Luo R, Sadakane K, Lam T-W (2015) MEGAHIT: an ultra-fast single-node solution for large and complex metagenomics assembly via succinct de Bruijn graph. Bioinformatics 31: 1674–1676. doi:10.1093/bioinformatics/btv033.

Li D, Luo R, Liu C-M, Leung C-M, Ting H-F, Sadakane K, Yamashita H, Lam T-W (2016) MEGAHIT v1. 0: a fast and scalable metagenome assembler driven by advanced methodologies and community practices. Methods 102: 3-11. 10.1016/j.ymeth.2016.02.020.

Meyer F, Fritz A, Deng Z-L, Koslicki D, Lesker TR, Gurevich A, Robertson G, Alser M, Antipov D, Beghini F, Bertrand D, Brito JJ, Brown CT, Buchmann J, Buluç A, Chen B, Chikhi R, Clausen PTLC, Cristian A, Dabrowski PW, Darling AE, Egan R, Eskin E, Georganas E, Goltsman E, Gray MA, Hansen LH, Hofmeyr S, Huang P, Irber L, Jia H, Jørgensen TS, Kieser SD, Klemetsen T, Kola A, Kolmogorov M, Korobeynikov A, Kwan J, LaPierre N, Lemaitre C, Li C, Limasset A, Malcher-Miranda F, Mangul S, Marcelino VR, Marchet C, Marijon P, Meleshko D, Mende DR, Milanese A, Nagarajan N, Nissen J, Nurk S, Oliker L, Paoli L, Peterlongo P, Piro VC, Porter JS, Rasmussen S, Rees ER, Reinert K, Renard B, Robertsen EM, Rosen GL, Ruscheweyh H-J, Sarwal V, Segata N, Seiler E, Shi L, Sun F, Sunagawa S, Sørensen SJ, Thomas A, Tong C, Trajkovski M, Tremblay J, Uritskiy G, Vicedomini R, Wang Z, Wang Z, Wang Z, Warren A, Willassen NP, Yelick K, You R, Zeller G, Zhao Z, Zhu S, Zhu J, Garrido-Oter R, Gastmeier P, Hacquard S, Häußler S, Khaledi A, Maechler F, Mesny F, Radutoiu S, Schulze-Lefert P, Smit N, Strowig T, Bremges A, Sczyrba A, McHardy AC (2022) Critical Assessment of Metagenome Interpretation: the second round of challenges. Nature Methods 19: 429–440. doi:10.1038/s41592-022-01431-4.

Mikheenko A, Prjibelski A, Saveliev V, Antipov D, Gurevich A (2018) Versatile genome assembly evaluation with QUAST-LG. Bioinformatics 34: i142–i150. doi:10.1093/bioinformatics/bty266.

Munk P, Brinch C, Møller FD, Petersen TN, Hendriksen RS, Seyfarth AM, Kjeldgaard JS, Svendsen CA, van Bunnik B, Berglund F, Bego A, Power P, Rees C, Lambrinidis D, Neilson EHJ, Gibb K, Coventry K, Collignon P, Cassar S, Allerberger F, Begum A, Hossain ZZ, Worrell C, Vandenberg O, Pieters I, Victorien DT, Gutierrez ADS, Soria F, Grujić VR, Mazalica N, Rahube TO, Tagliati CA, Rodrigues D, Oliveira G, de Souza LCR, Ivanov I, Juste BI, Oumar T, Sopheak T, Vuthy Y, Ngandjio A, Nzouankeu A, Olivier ZAAJ, Yost CK, Kumar P, Brar SK, Tabo D-A, Adell AD, Paredes-Osses E, Martinez MC, Cuadros-Orellana S, Ke C, Zheng H, Baisheng L, Lau LT, Chung T, Jiao X, Yu Y, JiaYong Z, Morales JFB, Valencia MF, Donado-Godoy P, Coulibaly KJ, Hrenovic J, Jergović M, Karpíšková R, Deogratias ZN, Elsborg B, Hansen LT, Jensen PE, Abouelnaga M, Salem MF, Koolmeister M, Legesse M, Eguale T, Heikinheimo A, Le Guyader S, Schaeffer J, Villacis JE, Sanneh B, Malania L, Nitsche A, Brinkmann A, Schubert S, Hesse S, Berendonk TU, Saba CKS, Mohammed J, Feglo PK, Banu RA, Kotzamanidis C, Lytras E, Lickes SA, Kocsis B, Solymosi N, Thorsteinsdottir TR, Hatha AM, Ballal M, Bangera SR, Fani F, Alebouyeh M, Morris D, O’Connor L, Cormican M, Moran-Gilad J, Battisti A, Diaconu EL, Corno G, Di Cesare A, Alba P, Hisatsune J, Yu L, Kuroda M, Sugai M, Kayama S, Shakenova Z, Kiiyukia C, Ng’eno E, Raka L, Jamil K, Fakhraldeen SA, Alaati T, Bērziņš A, Avsejenko J, Kokina K, Streikisa M, Bartkevics V, Matar GM, Daoud Z, Pereckienė A, Butrimaite-Ambrozeviciene C, Penny C, Bastaraud A, Rasolofoarison T, Collard J-M, Samison LH, Andrianarivelo MR, Banda DL, Amin A, Rajandas H, Parimannan S, Spiteri D, Haber MV, Santchurn SJ, Vujacic A, Djurovic D, Bouchrif B, Karraouan B, Vubil DC, Pal P, Schmitt H, van Passel M, Jeunen G-J, Gemmell N, Chambers ST, Mendoza FP, Huete-Pιrez J, Vilchez S, Ahmed AO, Adisa IR, Odetokun IA, Fashae K, Sørgaard A-M, Wester AL, Ryrfors P, Holmstad R, Mohsin M, Hasan R, Shakoor S, Gustafson NW, Schill CH, Rojas MLZ, Velasquez JE, Magtibay BB, Catangcatang K, Sibulo R, Yauce FC, Wasyl D, Manaia C, Rocha J, Martins J, Álvaro P, Di Yoong Wen D, Shin H, Hur H-G, Yoon S, Bosevska G, Kochubovski M, Cojocaru R, Burduniuc O, Hong P-Y, Perry MR, Gassama A, Radosavljevic V, Tay MYF, Zuniga-Montanez R, Wuertz S, Gavačová D, Pastuchová K, Truska P, Trkov M, Keddy K, Esterhuyse K, Song MJ, Quintela-Baluja M, Lopez MG, Cerdà-Cuéllar M, Perera RRDP, Bandara NKBKRGW, Premasiri HI, Pathirage S, Charlemagne K, Rutgersson C, Norrgren L, Örn S, Boss R, Van der Heijden T, Hong Y-P, Kumburu HH, Mdegela RH, Hounmanou YMG, Chonsin K, Suthienkul O, Thamlikitkul V, de Roda Husman AM, Bidjada B, Njanpop-Lafourcade B-M, Nikiema-Pessinaba SC, Levent B, Kurekci C, Ejobi F, Kalule JB, Thomsen J, Obaidi O, Jassim LM, Moore A, Leonard A, Graham DW, Bunce JT, Zhang L, Gaze WH, Lefor B, Capone D, Sozzi E, Brown J, Meschke JS, Sobsey MD, Davis M, Beck NK, Sukapanpatharam P, Truong P, Lilienthal R, Kang S, Wittum TE, Rigamonti N, Baklayan P, Van CD, Tran DMN, Do Phuc N, Kwenda G, Larsson DGJ, Koopmans M, Woolhouse M, Aarestrup FM, Global Sewage Surveillance C (2022) Genomic analysis of sewage from 101 countries reveals global landscape of antimicrobial resistance. Nature Communications 13: 7251. doi:10.1038/s41467-022-34312-7.

Murray CJ, Ikuta KS, Sharara F, Swetschinski L, Aguilar GR, Gray A, Han C, Bisignano C, Rao P, Wool E (2022) Global burden of bacterial antimicrobial resistance in 2019: a systematic analysis. The Lancet 399: 629–655. 10.1016/S0140-6736(21)02724-0.

Pruden A, Vikesland PJ, Davis BC, de Roda Husman AM (2021) Seizing the moment: now is the time for integrated global surveillance of antimicrobial resistance in wastewater environments. Current Opinion in Microbiology 64: 91–99. 10.1016/j.mib.2021.09.013.

Sczyrba A, Hofmann P, Belmann P, Koslicki D, Janssen S, Dröge J, Gregor I, Majda S, Fiedler J, Dahms E, Bremges A, Fritz A, Garrido-Oter R, Jørgensen TS, Shapiro N, Blood PD, Gurevich A, Bai Y, Turaev D, DeMaere MZ, Chikhi R, Nagarajan N, Quince C, Meyer F, Balvočiūtė M, Hansen LH, Sørensen SJ, Chia BKH, Denis B, Froula JL, Wang Z, Egan R, Don Kang D, Cook JJ, Deltel C, Beckstette M, Lemaitre C, Peterlongo P, Rizk G, Lavenier D, Wu Y-W, Singer SW, Jain C, Strous M, Klingenberg H, Meinicke P, Barton MD, Lingner T, Lin H-H, Liao Y-C, Silva GGZ, Cuevas DA, Edwards RA, Saha S, Piro VC, Renard BY, Pop M, Klenk H-P, Göker M, Kyrpides NC, Woyke T, Vorholt JA, Schulze-Lefert P, Rubin EM, Darling AE, Rattei T, McHardy AC (2017) Critical Assessment of Metagenome Interpretation—a benchmark of metagenomics software. Nature Methods 14: 1063–1071. doi:10.1038/nmeth.4458.

Su JQ, Cui L, Chen QL, An XL, Zhu YG (2017) Application of genomic technologies to measure and monitor antibiotic resistance in animals. Annals of the New York Academy of Sciences 1388: 121–135.

Villanueva RAM, Chen ZJ (2019) ggplot2: elegant graphics for data analysis. Taylor & Francis, pp. 10.1080/15366367.2019.1565254.

Vollmers J, Wiegand S, Kaster A-K (2017) Comparing and evaluating metagenome assembly tools from a microbiologist’s perspective-not only size matters! PloS one 12: e0169662. 10.1371/journal.pone.0169662.

Wang Z, Wang Y, Fuhrman JA, Sun F, Zhu S (2019) Assessment of metagenomic assemblers based on hybrid reads of real and simulated metagenomic sequences. Briefings in Bioinformatics 21: 777–790. doi:10.1093/bib/bbz025.

Wick RR, Schultz MB, Zobel J, Holt KE (2015) Bandage: interactive visualization of de novo genome assemblies. Bioinformatics 31: 3350–3352.

Xie H, Yang C, Sun Y, Igarashi Y, Jin T, Luo F (2020) PacBio long reads improve metagenomic assemblies, gene catalogs, and genome binning. Frontiers in Genetics 11: 516269. 10.3389/fgene.2020.516269.

Yi X, Liang J-L, Su J-Q, Jia P, Lu J-l, Zheng J, Wang Z, Feng S-w, Luo Z-h, Ai H-x, Liao B, Shu W-s, Li J-t, Zhu Y-G (2022) Globally distributed mining-impacted environments are underexplored hotspots of multidrug resistance genes. The ISME Journal 16: 2099–2113. doi:10.1038/s41396-022-01258-z.

Yorki S, Shea T, Cuomo CA, Walker BJ, LaRocque RC, Manson AL, Earl AM, Worby CJ (2023) Comparison of long- and short-read metagenomic assembly for low-abundance species and resistance genes. Briefings in Bioinformatics 24: doi:10.1093/bib/bbad050.

Zerbino DR, Birney E (2008) Velvet: algorithms for de novo short read assembly using de Bruijn graphs. Genome research 18: 821–829. 10.1101/gr.074492.107.

Zhang A-N, Gaston JM, Dai CL, Zhao S, Poyet M, Groussin M, Yin X, Li L-G, van Loosdrecht MCM, Topp E, Gillings MR, Hanage WP, Tiedje JM, Moniz K, Alm EJ, Zhang T (2021) An omics-based framework for assessing the health risk of antimicrobial resistance genes. Nature Communications 12: 4765. doi:10.1038/s41467-021-25096-3.

