## Supplementary Figures for "Metagenomic assemblies tend to break around antibiotic resistance genes"

Worst Median Best

A

| Statistics without reference | Velvet_0.5x_assembly | Ray_0.5x_assembly | SPAdes_0.5x_assembly | MEGAHIT_0.5x_assembly | metaSPAdes_0.5x_assembly | Trinity_0.5x_assembly |
|---|---|---|---|---|---|---|
| # contigs | 32 218 | 17 350 | 39 226 | 51 677 | 41 734 | 59 194 |
| Largest contig | 25 401 | 474 180 | 172 683 | 50 052 | 688 925 | 126 557 |
| Total length | 36 029 448 | 37 341 447 | 76 326 026 | 66 984 094 | 70 838 814 | 94 470 748 |
| Total length (>= 1000 bp) | 21 410 355 | 29 786 251 | 59 693 540 | 44 432 466 | 52 561 389 | 70 097 821 |
| Total length (>= 10000 bp) | 691 014 | 17 123 490 | 28 635 393 | 5 099 292 | 22 121 808 | 18 719 493 |
| Total length (>= 50000 bp) | 0 | 9 782 283 | 11 249 113 | 50 052 | 10 647 200 | 2 771 112 |
| **Mismatches** | | | | | | |
| # N's per 100 kbp | 0.06 | 0 | 0 | 0 | 0 | 0 |

| Statistics without reference | Velvet_1x_assembly | Ray_1x_assembly | SPAdes_1x_assembly | MEGAHIT_1x_assembly | metaSPAdes_1x_assembly | Trinity_1x_assembly |
|---|---|---|---|---|---|---|
| # contigs | 22 246 | 17 421 | 45 074 | 37 579 | 41 864 | 59 258 |
| Largest contig | 52 173 | 442 619 | 184 495 | 62 496 | 688 925 | 120 871 |
| Total length | 26 584 994 | 37 998 115 | 76 946 301 | 47 137 002 | 71 305 149 | 94 588 910 |
| Total length (>= 1000 bp) | 16 351 246 | 30 466 532 | 57 532 964 | 30 430 331 | 52 983 245 | 70 233 428 |
| Total length (>= 10000 bp) | 2 250 480 | 17 645 003 | 23 024 395 | 3 526 028 | 22 456 603 | 18 546 825 |
| Total length (>= 50000 bp) | 52 173 | 9 839 473 | 5 600 849 | 62 496 | 10 650 644 | 2 250 526 |
| **Mismatches** | | | | | | |
| # N's per 100 kbp | 2.18 | 0 | 0 | 0 | 0 | 0 |

| Statistics without reference | Velvet_5x_assembly | Ray_5x_assembly | SPAdes_5x_assembly | MEGAHIT_5x_assembly | metaSPAdes_5x_assembly | Trinity_5x_assembly |
|---|---|---|---|---|---|---|
| # contigs | 22 686 | 17 364 | 42 987 | 51 749 | 41 505 | 59 108 |
| Largest contig | 56 951 | 303 232 | 172 683 | 66 255 | 542 864 | 120 871 |
| Total length | 27 444 611 | 37 964 392 | 77 214 295 | 67 927 289 | 72 257 839 | 96 105 718 |
| Total length (>= 1000 bp) | 17 094 398 | 30 531 697 | 59 320 912 | 45 418 176 | 54 170 521 | 71 899 322 |
| Total length (>= 10000 bp) | 2 367 609 | 17 232 080 | 23 028 708 | 5 665 544 | 23 448 368 | 20 130 452 |
| Total length (>= 50000 bp) | 56 951 | 8 973 026 | 5 631 413 | 173 336 | 11 114 111 | 2 700 307 |
| **Mismatches** | | | | | | |
| # N's per 100 kbp | 1.89 | 0 | 0 | 0 | 0 | 0 |

| Statistics without reference | Velvet_10x_assembly | Ray_10x_assembly | SPAdes_10x_assembly | MEGAHIT_10x_assembly | metaSPAdes_10x_assembly | Trinity_10x_assembly |
|---|---|---|---|---|---|---|
| # contigs | 32 332 | 17 048 | 41 502 | 51 459 | 41 392 | 58 988 |
| Largest contig | 56 951 | 467 217 | 172 683 | 91 584 | 464 577 | 168 064 |
| Total length | 37 378 587 | 41 056 751 | 76 752 084 | 67 809 374 | 72 108 940 | 96 703 227 |
| Total length (>= 1000 bp) | 22 846 293 | 33 658 143 | 59 724 042 | 45 380 969 | 54 028 094 | 72 491 285 |
| Total length (>= 10000 bp) | 1 225 558 | 21 280 425 | 23 814 426 | 6 114 375 | 23 721 278 | 20 821 199 |
| Total length (>= 50000 bp) | 56 951 | 11 519 292 | 6 537 236 | 495 211 | 11 189 684 | 3 939 571 |
| **Mismatches** | | | | | | |
| # N's per 100 kbp | 0.08 | 0 | 0 | 0 | 0 | 0 |

B

| Statistics without reference | Velvet_assembly | Ray_assembly | SPAdes_assembly | MEGAHIT_assembly | metaSPAdes_assembly | Trinity_assembly |
|---|---|---|---|---|---|---|
| # contigs | 110 673 | 48 047 | 85 453 | 105 763 | 92 846 | 156 257 |
| Largest contig | 47 990 | 582 509 | 556 300 | 295 746 | 400 849 | 78 113 |
| Total length | 151 747 721 | 145 421 628 | 242 507 147 | 232 009 297 | 239 314 901 | 284 725 193 |
| Total length (>= 1000 bp) | 105 135 528 | 126 431 369 | 210 564 245 | 190 909 531 | 204 605 784 | 225 398 292 |
| Total length (>= 10000 bp) | 12 339 189 | 82 187 462 | 127 375 217 | 89 886 420 | 112 257 543 | 60 132 981 |
| Total length (>= 50000 bp) | 0 | 41 809 527 | 62 488 747 | 27 197 453 | 51 851 414 | 2 343 130 |
| **Mismatches** | | | | | | |
| # N's per 100 kbp | 0 | 0 | 0 | 0 | 0 | 0 |

**Figure S1 METAQUAST reports for assemblies using A) simulated short reads and B) using real data.**

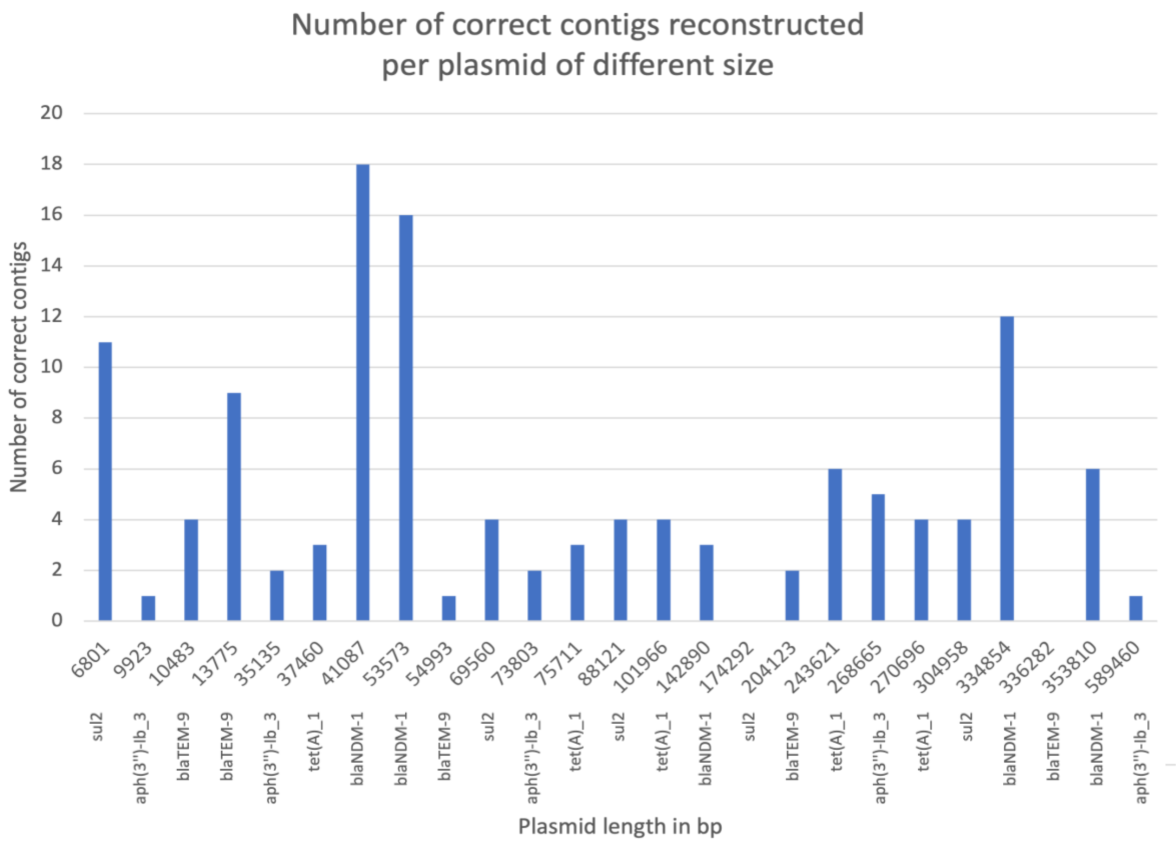


**Figure S2. Number of correct contigs per plasmid of different size.**

**Table S1. Correct genomic contexts assembled from Illumina short reads and compared to PacBio long reads as a reference.**

| Assembler | Number of unique contigs matching PacBio reads (98% identity, 100% coverage) | Number of different contexts the contigs match to | Average contig length, bp |
|---|---|---|---|
| Trinity | 10 | 10 | 2737 |
| SPAdes | 5 | 36 | 1555 |
| MEGAHIT | 5 | 30 | 1190 |
| metaSPAdes | 5 | 34 | 1061 |
| Velvet | 1 | 3 | 953 |
| TriMetAss | 1 | 5 | 652 |
| Ray | 3 | 26 | 968 |
