## Additional File 1 for "Metagenomic assemblies tend to break around antibiotic resistance genes"

**Simplified scenario with only two plasmids sharing the same ARG**
To test whether the reduced complexity of the simulated scenario can improve the assembly performance, instead of 25 plasmids we used only two. We performed this test with CP055250.1 (53573bp) and AP023079.1 (41087bp) both carrying *blaNDM-1* gene and CP064948.1 (589460bp) and CP039146.1 (73803bp) carrying *aph(3'')-Ib* gene. To create differential coverage we generated reads in proportions 2:1 for the first pair and 7:1 for the second pair of plasmids. The reads were generated and spiked into the real dataset as described for the simulated scenario in the materials and methods.

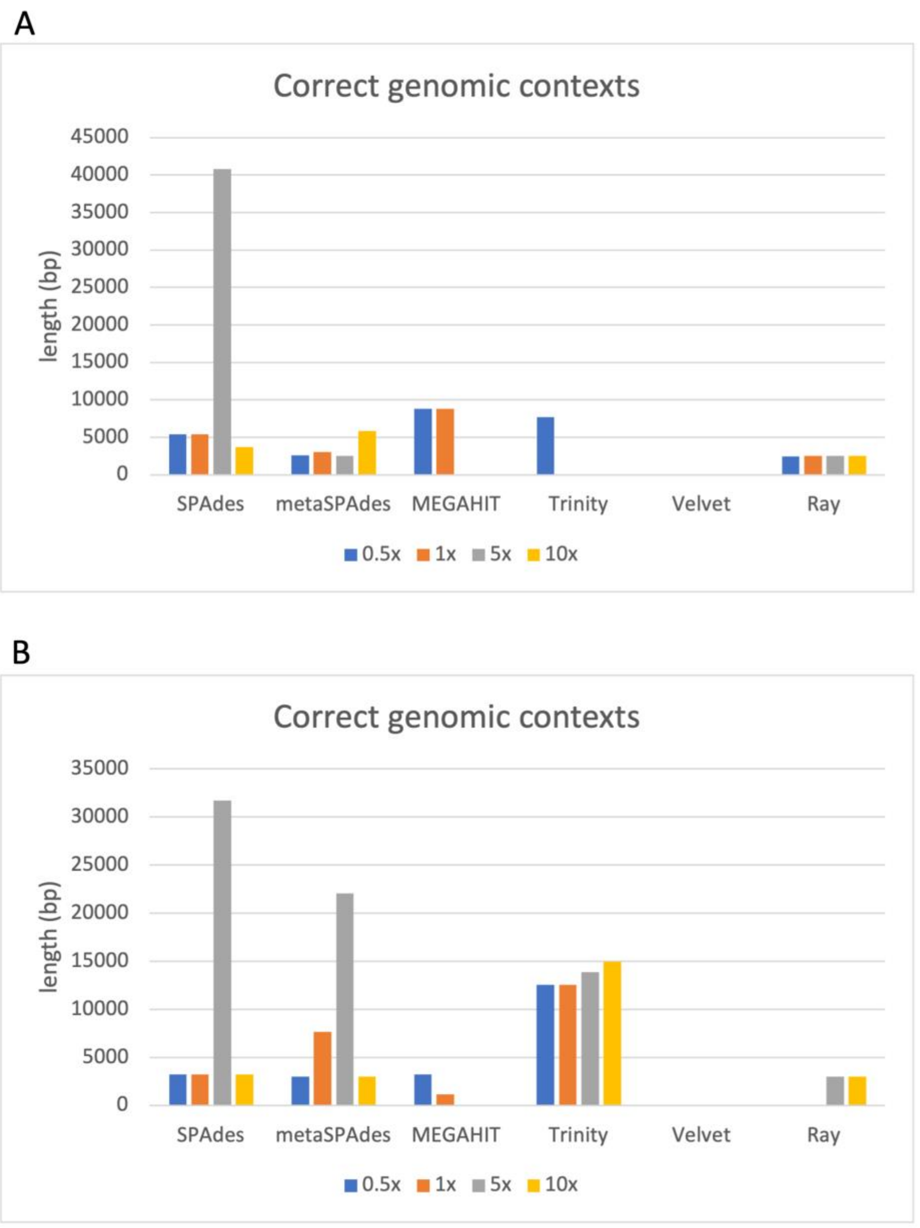


**Figure 1. Length of correct genomic contexts (containing full ARG sequence) assembled for the two plasmids carrying *aph(3'')-Ib* (A) and *blaNDM-1*(B).**

For the test with *aph(3'')-Ib* gene present on CP064948.1 and CP039146.1 plasmids, in all the assemblies only one correct genomic context from CP039146.1 plasmid was captured. This plasmid had the lowest generated coverage but much shorter length. Also, the *aph(3'')-Ib* gene on CP064948.1 is present in two copies and surrounded by several recombinases as well as the *aph(6)-I* gene in both cases, complicating the assembly (Figure 2). Despite that, assemblies with all four coverages produced at least two correct contigs of an average length of 3500 bp. In comparison, for the simulated scenario with 25 plasmids, only assemblies with 0.5x and 5x coverages produced correct contigs containing the *aph(3'')-Ib* gene but with an average length of 10,067 bp. In contrast, for Trinity only 5x and 10x assemblies generated correct contigs with an average length of 4491 bp, while in the simplified scenario only assembly with the lowest coverage produced one contig of 7681 bp. On the other hand, reduced complexity considerably improved performance of MEGAHIT (supported by results from Forouzan et al., 2018). This assembler did not produce any correct contigs in the simulated scenario with 25 plasmids but assembled correct contigs for the lowest coverage with average length of 8791 bp. In general, reduced complexity facilitated better performance for 5 out of 6 tools (in the scenario with 25 plasmids only 3 out of 6 tools produced correct contigs containing full ARGs).

As mentioned before, not all of the genes were equally easy to assemble. The assembly results from two plasmids carrying *blaNDM* were largely similar to the test with *aph(3'')-Ib* (Figure 1B). However, in comparison to the other tools Trinity showed considerable improvement. It produced correct contigs for all four assemblies of different coverages, and those contigs were on average three times longer than in the test with *aph(3'')-Ib* gene. This suggests that *blaNDM* is existing in contexts that are much easier to piece together. In addition, in two plasmids scenario Trinity also produced contigs two times longer than in the complex scenario with 25 plasmids.

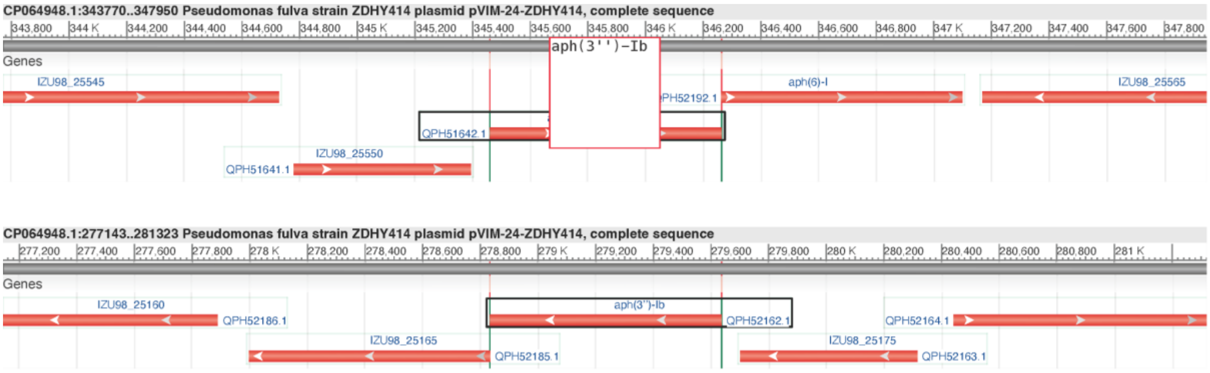


**Figure 2. Genomic context of *aph3''-Ib* gene on CP064948.1 plasmid.**
